# Leveraging whole genomes, mitochondrial DNA, and haploblocks to decipher complex demographic histories: an example from a broadly admixed arctic fish

**DOI:** 10.1101/2024.12.11.628006

**Authors:** Xavier Dallaire, Eric Normandeau, Thomas Brazier, Les Harris, Michael M. Hansen Michael, Claire Mérot, Jean-Sébastien Moore

## Abstract

The study of phylogeography has transitioned from mitochondrial haplotypes to genome-wide analyses, blurring the line between this field and population genomics. Whole-genome sequencing offers the opportunity to join use both and provides the density of markers necessary to investigate genetic linkage and recombination along the genome. This facilitates the unraveling of complex demographic histories of admixture between divergent lineages, as is often the case in species evolving in recently deglaciated habitats. In this study, we sequenced 1120 Arctic Char genomes from 33 populations across Canada and Western Greenland to characterize patterns of genetic variation and diversity, and how they are shaped by hybridization between the Arctic and Atlantic glacial lineages. Several lines of evidence supported mito-nuclear discordance in lineage distribution, with all Canadian populations under the 66^th^ parallel being characterized by introgression from the Atlantic lineage, leading to higher nuclear genetic diversity. By scanning the genome using local PCAs, we identified putative low-recombining haploblocks as local ancestry tracts from either lineage and described the impacts of recombination on the introgression landscape in admixed populations. Finally, we inferred conflicting origins of recolonization using whole genomes vs. ancestry tracts for the Arctic lineage, suggesting that haplotypes sheltered from introgression by low recombination could enlighten complex post-glacial histories. Overall, we argue that Whole-Genome Sequencing, even at low depths of coverage, provides a versatile approach to the study of phylogeographic dynamics.

## Introduction

Understanding a species’ evolutionary and demographic history is central to interpreting the processes that produced its contemporary genetic diversity and population structure. Since its inception, the field of phylogeography has aimed to interrogate the micro-evolutionary processes responsible for the geographical distribution of genetic lineages [1]. For two decades, phylogeography was dominated by the study of gene trees based on mitochondrial DNA (mtDNA) sequences because of their easy sequencing, short length, and simple uniparental inheritance without recombination [1,2]. With the advent of high-throughput sequencing, phylogeographic studies now commonly use thousands of nuclear DNA (nuDNA) markers such as Single Nucleotide Polymorphisms (SNPs). Frequent cases of discordance between mitochondrial and nuclear genetic variation have highlighted the limitations of mtDNA [3–5], e.g. the effect of sex-biased migration on genealogies for these matrilineally inherited markers. Despite these limitations, mtDNA remains a valuable tool in phylogeographic studies: being haploid, non-recombining, and having a lower effective population size, they provide a rich source of information independent from the nuclear genome. Used in conjunction with nuclear markers, they offer a more comprehensive understanding of species’ evolutionary histories and genetic diversity.

The integration of genome-wide nuclear markers into phylogeography has also blurred the distinctions between this field and population genetics [2]. Population genomics continues to rely heavily on averaging the genome-wide genetic variation of SNPs, for example, as a means to investigate population structure, while ignoring fluctuations in genetic variation at a finer genomic scale [6,7]. However, these “vertical” approaches stacking information from individual genetic markers across individuals usually ignore or purposely try to reduce the impacts of linkage disequilibrium, neglecting the “horizontal” signal contained in haplotypes [8]. Haplotypes are usually defined as DNA sequences where alleles for multiple markers are inherited as a block. Unlike SNP-based approaches focusing on mutations and assuming independence between markers, haplotypes carry information about recombination events and have the potential to increase accuracy in the inference of selection, gene flow, and population structure [9,10]. In particular, long haplotypes, sometimes called haploblocks, may be maintained in low recombination regions or within structural rearrangements, providing information about deeper coalescent times than short haplotypes within high recombination regions [11].

Phylogeographic studies are particularly relevant for arctic species, as their habitats and ranges were among the most impacted by the glacial cycles of the Pleistocene. As demographic bottlenecks and extended periods in allopatry lead to the accumulation of divergence between populations that survived in distinct glacial refugia [12–14], post-glacial recolonizations are commonly the stage for secondary contact between previously isolated lineages [15,16]. These cases of hybridization between lineages lead to an admixture of genetic backgrounds, producing a mosaic of local ancestry tracts that tend to shorten through the recombination of chromosomes over generations of backcrossing [8,17]. Whole-genome averaging approaches can struggle to properly identify admixture at a relatively recent glacial timescale, as isolation-by-distance can produce similar genomic signatures [18]. Therefore, integrating multiple aspects of the genome (e.g., mtDNA and haplotype information) may help distinguish between true admixture events and patterns that arise due to gradual changes in genetic variation over geographical space.

Arctic char (*Salvelinus alpinus*) is a widespread circumpolar species with the northernmost distribution observed in freshwater or diadromous fishes, most of which coincides with regions covered by ice sheets at the last glacial maximum, around 18,000 years ago [15,19]. Arctic Char has been called “the most diverse vertebrate on earth” [20] with many localities harboring morphs diverging in size, diet, habitat choice, phenology, migratory tactic, or life trait history traits (Reist et al. 2013). Some authors recognize dozens of subspecies, and five main glacial lineages have been defined using mtDNA [21,22]. However, these fail to offer a perfectly discrete classification of wild populations around the globe, as post-glacial secondary contact zones have been described between most pairs of neighboring mtDNA lineages [23–26]. With their unique blend of moderate philopatry, higher dispersal rate than other anadromous salmonids [27], and range spanning the most recently deglaciated region around the globe, Arctic Char offers unique opportunities for the study of the distribution of genetic diversity in post-recolonization populations. Notably, the origin of the post-glacial recolonization of the ‘Arctic’ mtDNA lineage, presently found throughout most of the Canadian Arctic, has remained cryptic, while both Beringia and the Arctic Archipelago have been suggested as potential refugia [22].

In this study, we used low-coverage whole-genome sequencing to investigate the recolonization of the recently deglaciated Canadian Arctic by several lineages of Arctic Char and their putative hybridization. We sequenced 1,120 Arctic Char genomes (Fig. 1a; average coverage = 2X, with 100 samples in 5 populations further sequenced at a depth of 8X) from 33 river systems across the 4,000 km wide range of the previously defined Arctic mitochondrial lineage in northern Canada and western Greenland to characterize patterns of genetic variation, diversity, and admixture with the neighboring Atlantic lineage. We combined information from both mitochondrial and nuclear genomes including mtDNA haplotype networks, genome-wide admixture and ancestry, low-recombining haploblocks, and demographic inference to resolve the history of the secondary contact between those glacial lineages. Finally, to highlight the complexity of this post-glacial evolutionary history, we aimed to infer the origin of recolonization of the Arctic lineage by comparing derived allele distributions across the genome and inside Arctic haploblocks potentially carrying different non-recombining ancestry.

**Figure 1:**
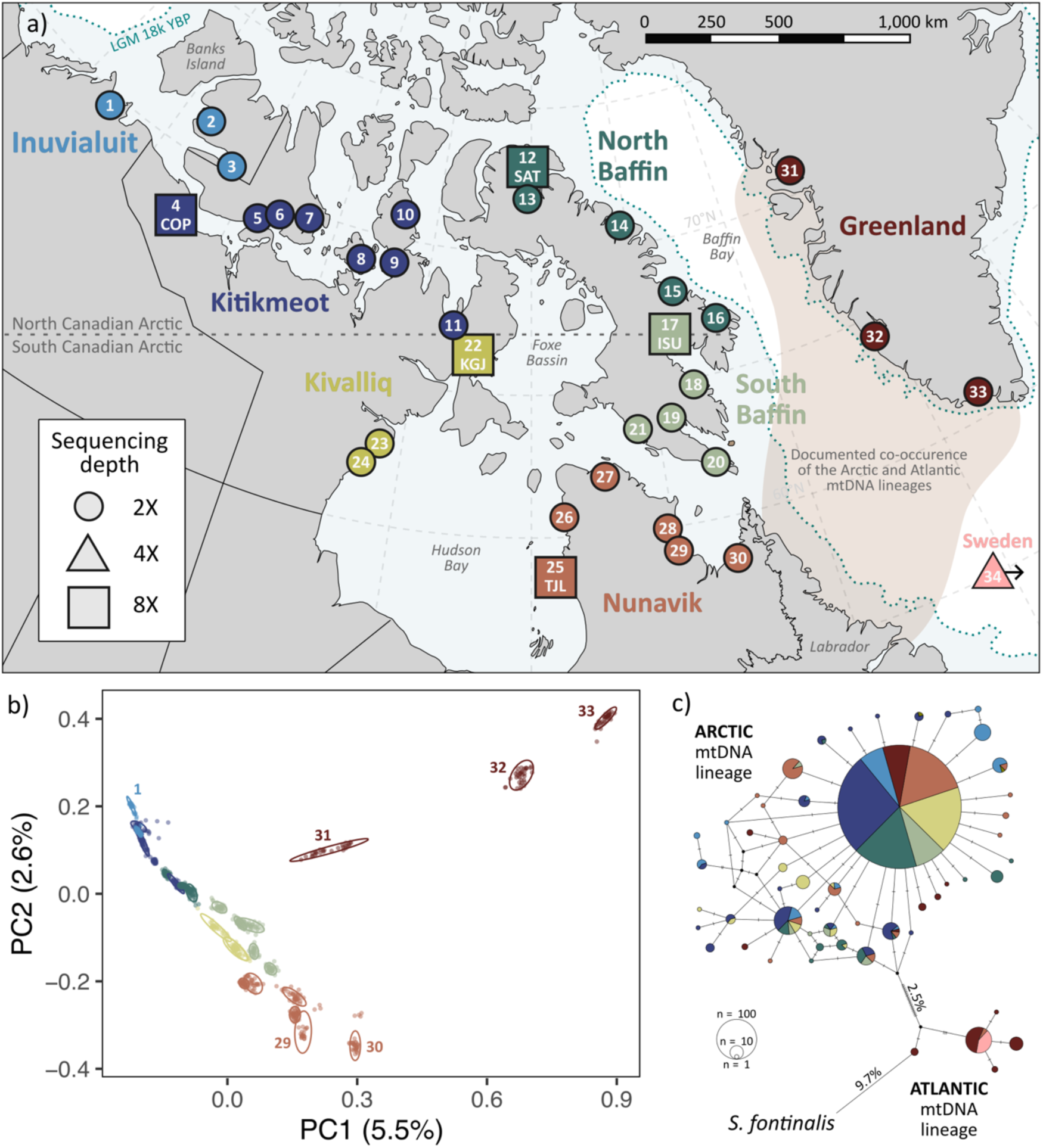
A highly structured distribution of genetic variation across the Arctic. a) Sampling sites for 1140 Arctic Chars (Salvelinus alpinus) sequenced at a coverage of 2X (circle), 5X (triangle), or 8X (square). Colors indicate sampling regions, and the same color coding is used in all figure panels. Dotted blue lines delimit the ice margin at the last glacial maximum (18 ka, Dalton et al. 2020). b) Principal components analysis based on 396,113 independent SNPs. 95% confidence interval ellipses are drawn around each sampling site. c) Median-joining haplotype network for the mitochondrial D-loop control region (993 bp), showing divergence between the Arctic (left) and Atlantic (right) mitochondrial lineages. For each haplotype shared by n>1 individual, colored sections of the pie chart indicate the proportional representation of individuals from each region.

## Results

### Distribution of nuclear genetic variation across the North American Arctic

Nuclear genome-wide variation at 5.98 million SNPs (average coverage = 2X, Fig. S1) in 1,120 anadromous Arctic Char from 33 Canadian and Greenlandic rivers (Fig. 1a, Table S1) displayed a strong substructure associated with rivers, which grouped into two major genetic clusters corresponding to Northern and Southern areas. Genome-wide pairwise F_ST_ ranged from 0.012 to 0.381 among Canadian rivers, while comparisons including Greenlandic sites reached up to F_ST_ = 0.632 (pop. 1 and 33) (Fig 2a). The general structure associated with longitude was visible on the first axis of a principal components analysis (Fig 1b), explaining 63.9% and 5.5% of the genome-wide variance, respectively before and after stringent LD-pruning (375,236 SNPs). As expected for homing salmonids, each river system appeared genetically differentiated in an NGSadmix analysis (Fig. S2).

**Figure 2:**
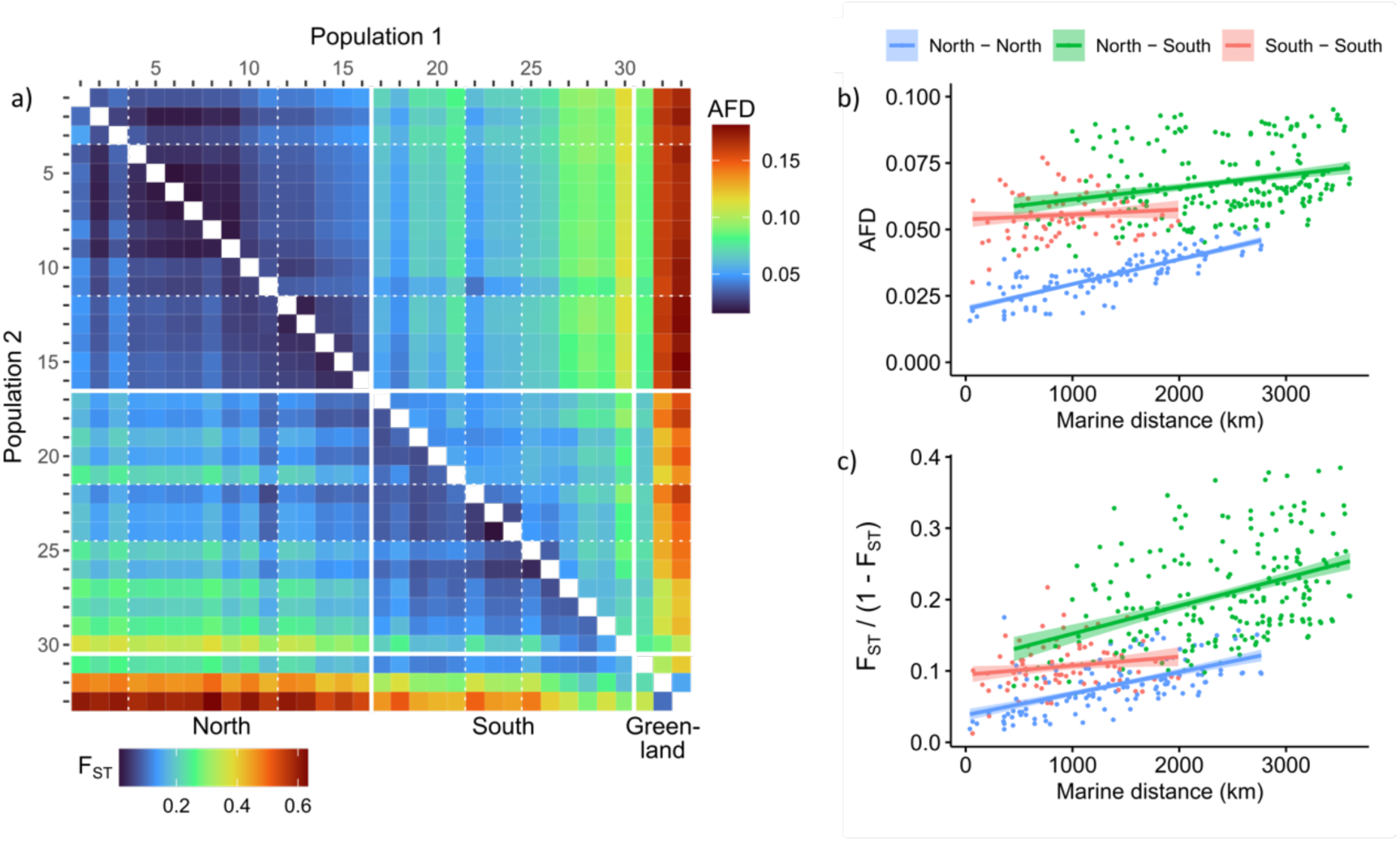
A North-South divide in genetic variation. a) Pairwise genetic differentiation among populations estimated using allele frequency difference (AFD, above diagonal) and genome-wide weighted FST (below diagonal). Greenlandic and Canadian populations above and below the 66th parallel are divided by solid white lines, and dotted lines divide regions defined in Figure 1a. b-c) Isolation-by-distance as estimated by the linear regression between genetic (either b) AFD or c) linearized FST) and marine distance between populations 1-29. Population pairs were categorized based on their inclusion in the North (blue, pop. 1-16) or South (red, pop. 17-29) group, or including a Northern and a Southern population (green).

Pairwise differentiation among populations, both based on the F_ST_ index and allele frequency differences (AFD), highlighted a genetic divide between the northern (pop. 1-16) and southern (pop. 17-30) Canadian Arctic around the 67^th^ parallel (Fig. 2a). While populations from both regions exhibited patterns of isolation-by-distance (IBD), southern populations were more differentiated at equal marine distances than northern populations (Fig. 2b-c). Greenlandic populations and pop. 30 (south-eastern Nunavik) were comparatively more differentiated than other populations and did not fit the Canadian IBD patterns. Intra-population genetic diversity, as measured by nucleotide diversity (ϴ_π_), was also higher in southern (median widowed per-site ϴ_π_ = 1.13-2.77 x 10^-3^) than in northern (ϴ_π_ = 0.72-2.09 x 10^-3^) Canadian populations (Fig. 3d).

**Figure 3:**
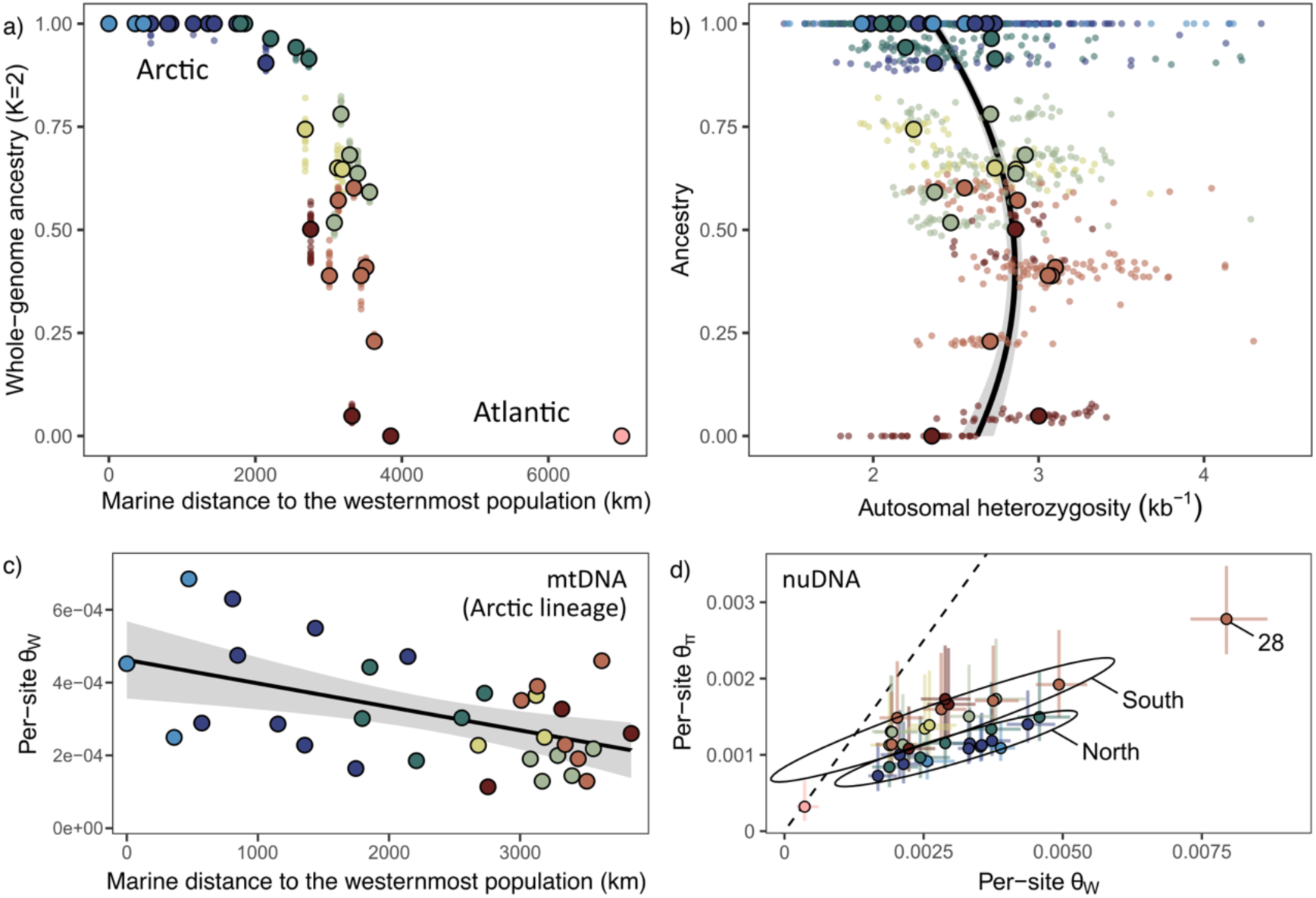
Higher nuclear diversity in admixed populations. a) Genome-wide Arctic ancestry, estimated by admixture proportion in NGSadmix (K = 2), of individuals (smaller points) and sampling sites (mean ancestry, circled points) along a spatial gradient from Inuvialuit (left) to Sweden (right). b) Individual autosomal heterozygosity (heterozygote sites per kb) is higher in the secondary contact zone (intermediate ancestry), as visualized by a fitted quadratic equation. c) Per-site Watterson’s estimator (ϴ_W_) in mitochondrial haplotypes of the Arctic lineage by population. d) Per-site nucleotide diversity (ϴ_π_) and Watterson’s estimator by population, estimated in 100 kb windows along the nuclear genome. The median (point), 25^th^, and 75^th^ percentiles (error bars) are shown. The Northern (pop. 1-16) and Southern (pop. 17-30) Canadian populations are highlighted by 95% confidence interval ellipses and a dotted line follows ϴ_π_ = ϴ_W_ (Tajima’s D = 0).

### Admixture between lineages is a major determinant of variation in southern populations

Several lines of evidence support the hypothesis that the large intra- and inter-population diversity described earlier in the Southern Canadian Arctic and Greenland is due to admixture between the Arctic and Atlantic glacial lineages.

First, we realigned and genotyped data from 20 Arctic Char collected in a Swedish population (pop. 34, Saha et al. 2024; depth of coverage = 5X) to serve as an outgroup from the Atlantic lineage and conducted an admixture analysis (K=2, NGSadmix). Southern Canadian and north-western Greenlandic populations had mixed genome-wide ancestry for the two groups, distributed along a cline-like northwest-to-southeast gradient, suggesting admixture between the Arctic (i.e., northern Canadian populations) and Atlantic (i.e., Swedish outgroup) lineages (Fig. 3a). Within population variance in proportions of Atlantic ancestry was low (Fig. 3a), and putatively admixed populations had higher observed heterozygosity (Fig. 3b, Fig. S3) and nucleotide diversity (Fig. 3d), supporting the hypothesis of secondary contact between the Arctic and Atlantic lineages.

Second, to further test the hypothesis of admixture between lineages in the southern part of the study area, we computed *f_4_* statistics [28,29] using a Dolly Varden (*Salvelinus malma*) population from Yukon, Canada, as an outgroup. This test provided support for introgression from the Atlantic lineage (Sweden) toward the southern Arctic populations (Fig. 4c). Finally, using demographic modeling performed with fastsimcoal, with the Swedish population and 8X data for COP (pop. 4) and TJL (pop. 25), a model including ancient gene flow between the Atlantic lineage and the southern Canadian Arctic was better supported than models with no or more recent gene flow (Fig. 4a-b; see demographic parameter estimation in Fig. S4).

**Figure 4:**
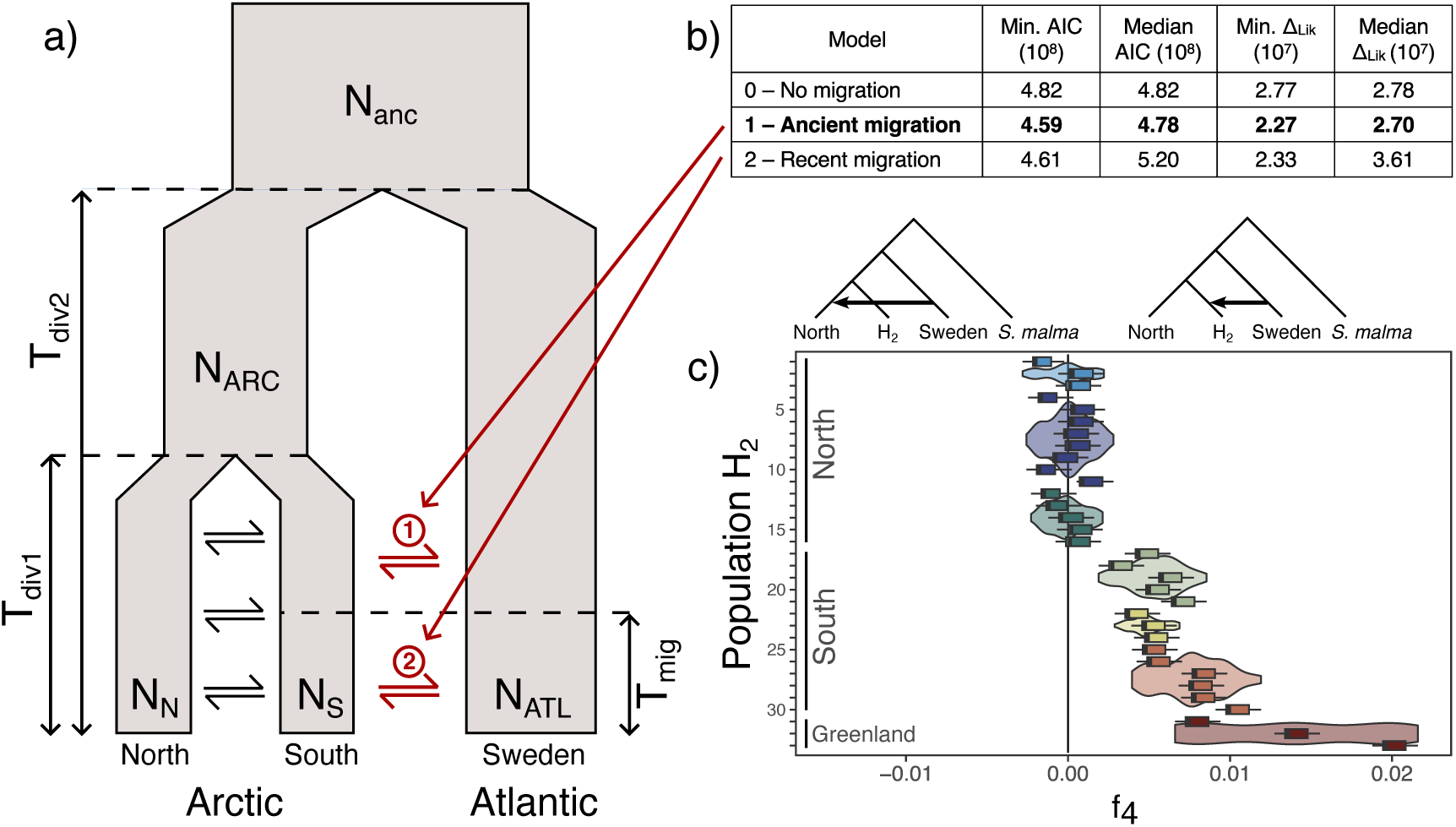
Evidence for gene flow between the Atlantic and South Arctic. a) Simplified demographic model showing ancient divergence between the Arctic and Atlantic, followed by isolation with migration of the North (COP, pop. 4) and South (TJL, pop. 24) Arctic. b) Support (AIC and Δ_Lik_, i.e. difference between observed and predicted likelihood of the SFS) for alternative fastsimcoal models (100 runs per model) for 0) absence of gene flow, 1) ancient or 2) recent gene flow between the South Arctic and Atlantic lineage (represented by the Swedish population), as highlighted in red in panel a). c) f_4_ test for introgression from the Atlantic lineage (represented by the Swedish population), where a negative f_4_ statistic supports introgression in population H_1_ (only tests with populations from northern regions are shown) and a positive f_4_ supports introgression in population H_2_ (detailed on the y-axis). Similar lcWGS data from Dolly Varden (Salvelinus malma) was used as an outgroup.

### Mito-nuclear discordance in admixture and diversity patterns

To leverage an independent source of historical demographic information and to compare our results with previous studies, we extracted mitochondrial D-loop sequences (993 bp) from whole-genome data (Fig 1c). Consistent with previously published data based on mtDNA, but contrary to our results using the nuclear genome, the Arctic lineage dominates in populations of the entire Arctic archipelago outside Labrador [22,23,26], with only one Canadian sample from the present study carrying an Atlantic D-loop haplotype (in pop. 28). In contrast, the Atlantic and Arctic mitochondrial lineages co-occurred in western Greenland (Jacobsen et al. 2021). As previously observed, a single mitochondrial haplotype dominated the Arctic lineage (77.5% of samples shared the most common D-loop haplotype). In Greenlandic populations, the individuals carrying haplotypes from either mtDNA lineages did not differ in their nuclear genome ancestry level inferred with NGSadmix (t-test per population: p > 0.05).

Whole-mitogenome genetic diversity inside the Arctic mtDNA lineage also displayed different trends compared to the nuclear genome. While nuclear diversity was the highest in the admixed populations of the South Arctic and Greenland, both mitochondrial ϴ_W_ and haplotype diversity were higher near the westernmost population than in the southeastern part of the study area (Fig. 3c, Fig. S5a,d). Mitochondrial nucleotide diversity (ϴ_π_) and Tajima’s D were lower at intermediate distances from the westernmost population, i.e. the center of the Arctic mtDNA lineage range. Finally, the frequency of the most frequent whole-mitogenome haplotype, present in 34% of all Arctic lineage samples, was lower in populations at both extremities of the range of the lineage, i.e. in Inuvialuit and Greenland (Fig. S5e).

### Heterogeneity in the introgression along the genome

To investigate the fine-scale genomic landscape of admixture between the Arctic and Atlantic lineages, we scanned the genome for distinct patterns of genetic variation and detected putative local ancestry tracts for both lineages.

We used local Principal Component Analyses (PCA) and multidimensional scaling (MDS) in 100 SNPs-wide windows to identify genomic regions displaying patterns of genetic variation that were significantly different from the chromosome-wide population structure. We retained 117 independent regions that contained at least 10 MDS outlier windows (e.g. Fig. 5a, see all outlier regions in Fig S6-7). These regions ranged from 98 kb to 5.26 Mb in length (median = 825 kb, total = 133 Mb). Of these regions, we visually identified 46 that showed three clear clusters on a first PC explaining most of the variance (e.g., Fig. 5b). We interpret these regions as putative haplotype blocks, or haploblocks, i.e. regions of low recombination where polymorphic sites are inherited in high linkage, creating groups of haplotypes, or haplogroups. As such, the three observed clusters correspond to the two homozygotes and the heterozygote for these haplogroups. Using populational-scale recombination maps based on 8X data (Fig. S8), we confirmed that the putative haploblocks tended to have lower recombination rates than the chromosome-wide average (Fig. S9), further supporting this interpretation.

**Figure 5:**
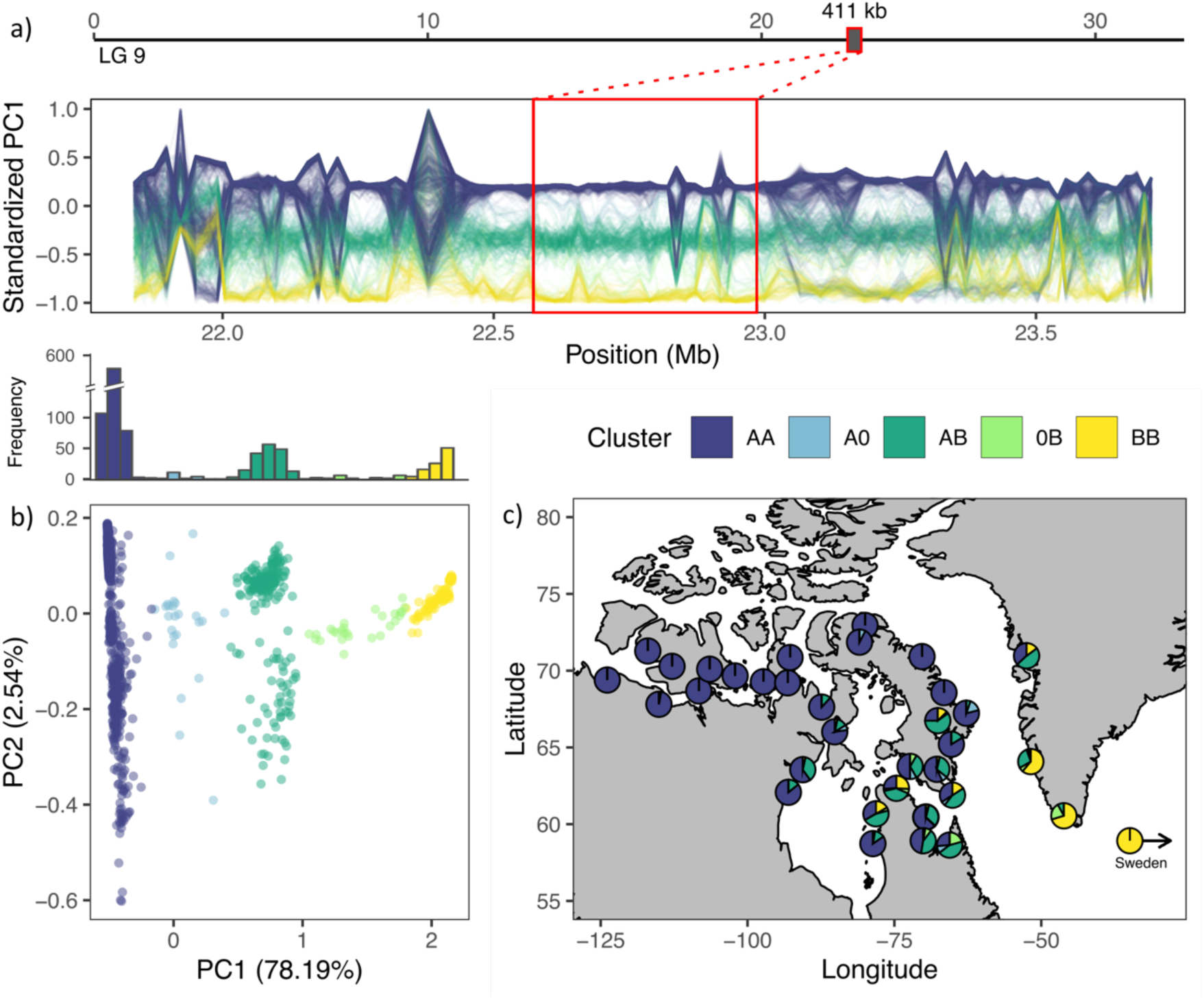
Local ancestry inferred through local PCA. Representative case of a low-recombining diverged putative haploblock corresponding to a local PCA outlier region (win8) on LG9 (NC_036849.1). a) Position of the MDS outlier region (red) on the linkage group (above) and standardized PC1 coordinates in PCAs for successive non-overlapping 100-SNP windows inside and around the outlier region (below). Each sample is represented by a line colored according to their haplogroup inferred from panel b). b). PCA for SNPs in the outlier region where the three major clusters found on PC1 are attributed to the 2 homozygotes (AA, blue, left; BB, yellow, right) and their heterozygote (AB, green, middle). Intermediate genotypes (A0, light blue; 0B, light green) were assigned to individuals outside of the cluster distributions. The histogram (above) represents the density of individuals along the first axis of the PCA. c) Proportion of inferred haplogroups for the win8 haploblock across sampling sites.

Most of these putative haploblocks corresponded to islands of differentiation between the Arctic and Atlantic lineages. Twenty-three haploblocks (median length = 783 kb, total = 20 Mb) were differentially fixed between the north-western Canadian Arctic (Inuvialuit, Kitikmeot; henceforth identified as haplogroup ‘A’) and Sweden (haplogroup ‘B’) while showing heterozygotes in southern populations of the Canadian Arctic (e.g., Fig. 5c), suggesting that they may be tracts of different ancestry that are yet unbroken by recombination. The individual distribution of heterozygotes differed between haploblocks, supporting their independent genealogy, but haplogroup frequencies by population were correlated with the Arctic-Atlantic whole-genome ancestry inferred earlier (NGSadmix, K = 2). Such correlations further support a scenario of secondary contact and introgression at the nuclear level by revealing blocks of linked polymorphic markers that appear to be inherited from ancestral lineages, after their divergence in allopatry during past glacial periods.

Interestingly, two other MDS outliers (e.g., win21 and win58 in Fig. S7) also featured three distinct genotypes on PC1, but their spatial distribution did not match the Arctic-Atlantic admixture gradient. Those could be associated with recent and polymorphic structural variants suppressing recombination such as chromosomal inversions, allowing the accumulation of genetic divergence in sympatry.

### Inference of origin of recolonization

Directionality analysis conducted on either the whole genome or restricted to variation inside Arctic haplotypes supported conflicting recolonization histories, respectively from the east and west of the lineage range. Based on genome-wide data, we inferred an origin of recolonization around the southeast of the Canadian Arctic by comparing the spatial distribution of genome-wide derived alleles in Canadian populations through the directionality index (*ψ*) and Time Difference of Arrival (TDoA) (Fig. 6a). This is accomplished by testing every potential origin (cells on a spatial raster) for a positive correlation between population-pairwise *ψ* (i.e., difference in derived allele frequencies) and geographical distance from the origin (maximum R^2^ = 0.638 before rescaling). This inference is nevertheless to be taken with caution because Canadian populations deviate from the single-origin assumption of TDoA analysis and are precisely characterized by extensive introgression in the southeast.

**Figure 6:**
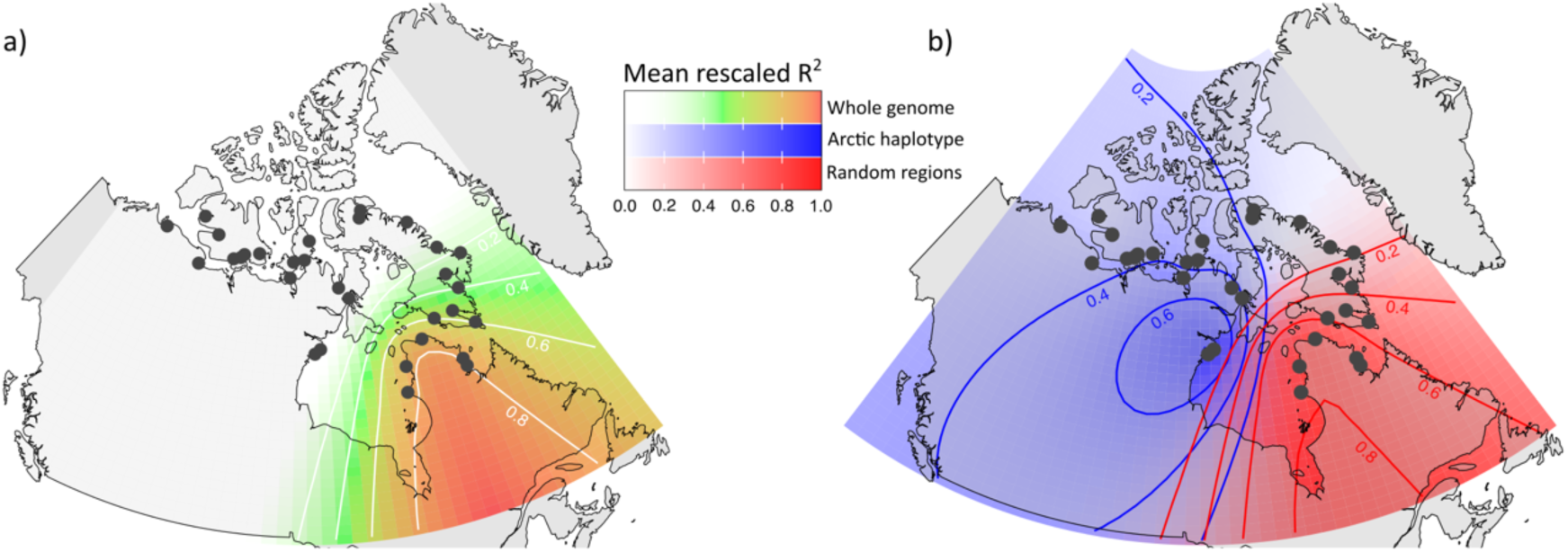
Genome-wide and Arctic lineage-specific inference of the origin of recolonization. a) Time Difference of Arrival (TDoA) analysis for inference of the origin of recolonization, where red color indicates a higher correlation (R^2^) between distance from the putative origin and whole-genome directionality index (ψ). b) TDoA for ψ calculated in Arctic haplogroup homozygotes (10 individuals per population) in 23 putative local ancestry tracks (total length = 20 Mb). The mean rescaled R^2^ for 100 replicate runs is shown in blue, and the mean rescaled R^2^ for 100 runs on 23 random genomic regions (total length = 20 Mb) is shown in red.

To confirm the directionality analysis on a putatively non-introgressed fraction of the genome, we leveraged the previously identified local ancestry tracts to compute *ψ* on derived allele frequencies in homozygotes for the Arctic lineage-specific alleles (e.g. haplogroup A in Fig. 5) of these local ancestry tracts (n = 23, total length = 20 Mb). Despite a reduced genomic size (1.3 % of the total genome), the resulting 100 replicate runs of TDoA weakly supported an origin of recolonization in the western part of the sampling area (maximum R^2^ ranging from 0.007 to 0.107, Fig 6b, Fig S10a), contrary to the eastern origin of recolonization inferred using genome-wide data.

To control for the small number of non-recombining haploblocks and the difficulty in precisely delimiting these putative local ancestry tracts, we ran the same analyses on random genomic regions of equivalent lengths. These analyses supported a south-eastern origin of recolonization, which was highly congruent with the whole-genome TDoA (maximum R^2^: 0.199 – 0.483, Fig. 6b, Fig. S10b). This suggests that, although our current prediction power is low, genetic diversity inside local ancestry tracts could carry distinct lineage-specific information that can be leveraged to elucidate more complex evolutionary histories.

## Discussion

It has long been recognized that making phylogeographic inferences based on mitochondria alone can lead to biased interpretations [30]. However, the non-recombining nature of mtDNA can offer advantages in specific cases since it can ‘record’ past events that are not blurred by generations of hybridization and admixture. At the same time, vast amounts of nuclear data are sometimes insufficient in revealing historical patterns if averaged over the genome, and evolutionary inferences could benefit from identifying haploblocks that retain historical signals [8,10]. In this study, we sought to maximize the use of whole-genome sequencing data by integrating mitochondrial and nuclear analyses, both at the genome-wide and haplotypic levels, to assess the extent of admixture between two glacial lineages of an arctic fish. Intraspecific hybridization between glacial lineages and the resulting increased nuclear diversity observed in southern populations was a dominant feature of the study system. Such introgression across lineages obscured much of the genomic signal for post-glacial recolonization from either single lineage. We scanned the genome for regions of distinct genetic variation and identified putative local tracts of conserved Arctic or Atlantic ancestry. By comparing genetic variation within Arctic tracts to the whole genome, we uncovered a putative cryptic glacial refugium for the lineage, despite a heavily admixed genetic background.

### Extensive admixture between lineages and mito-nuclear discordance

In Arctic Char, we show evidence for the genetic admixture of the Arctic and Atlantic lineages in most of Arctic Canada under the 66^th^ parallel. This is in stark contrast with the previously described distribution of mitochondrial haplotypes, with observed co-occurrence zones of both lineages restricted to Labrador [26] and western Greenland [25]. While previous phylogeographic work on the species hinted at the introgression from the Atlantic lineage reaching further west at the nuclear level [22,23], here we inferred the presence of Atlantic nuclear alleles thousands of kilometers away from the range limit of the Atlantic mitochondrial lineage, both using f_4_ tests and indirect detection of lineage-divergent haploblocks.

This kind of mito-nuclear discordance has been observed in numerous natural systems [4] and could be explained by multiple mechanisms. First, as non-recombinant haploid sequences, mitogenomes have a population effective size (Ne) that is ¼ of their diploid nuclear counterparts [31] and are thus more impacted by genetic drift. Alternatively, sex-biased dispersion has been suggested in Arctic Char with males appearing more mobile than females [32,33]. As mitochondrial genomes are matrilineally inherited, this could contribute to a discordance in the distribution of nuclear DNA and mitochondrial DNA variation.

### Local PCA reveals more than just inversions

Short-read whole-genome sequencing has become increasingly accessible in non-model organisms. New analytical frameworks allow for the study of population genetics questions even with low individual depth of coverage, which has opened the door for whole-genome studies of unprecedented sampling sizes [34]. However, phasing (i.e. haplotype estimation) might be impossible in these designs without a reference panel [35], as haploblocks are most reliably defined and delimited using long-or linked-read data [36]. It has been suggested that, when they are large and divergent enough (e.g., in regions of low recombination), haploblocks might also be indirectly detectable through genome scans for distinct patterns of genetic variation [37]. One such method is the use of local PCAs, an approach that has risen as a popular approach to detect large recombination-suppressing structural variants (e.g., inversions) [38]. The typical signal expected around non-recombining haploblocks is a PCA forming three distinct clusters on the first axes (generally PC1), with the middle cluster carrying more heterozygous sites than the two others, and higher linkage disequilibrium among clusters than inside each one [38].

In this study, we identified local PCA outliers by performing multidimensional scaling (MDS) at the chromosomal level that allowed for the detection of relatively short (250 kb to 4 Mb) candidate regions on every chromosome. We analyzed the five first MDS axes, but Fig. S7 suggests that many additional outlier regions could be identified on subsequent axes. PC1 shows the expected three-cluster pattern indicative of haploblocks, i.e. the two homozygotes and the heterozygote for a region of low recombination, and inversions may underlie some of those haploblocks. For example, one of our MDS outlier regions indicative of a haploblock (win16) was located on the distal end of LG12, which was recently identified as a putative inversion in Nunavut Arctic Char [39]. However, we observed that the clustering in three groups was far from perfect in most of the haploblocks, with many intermediate individuals (e.g. Fig. 5B). This suggests a suppression of recombination less severe than what has been observed in recent publications investigating inversions [38,40]. Intriguingly, some candidate haploblocks (including win16) showed additional structure inside an inferred homozygote group on the second axis of the PCA, resulting in a triangular pattern when considering PC1 and 2 (e.g., win50, win89, win91, and win109 in Fig. S7), which may result from nested inversions or regions of low recombination maintaining more than two distinct groups of haplotypes. Therefore, overall, we proposed (and confirmed for some) that those regions are characterized by low recombination but do not further speculate with the present data about the precise mechanism protecting the observed haploblocks from recombination.

More interesting is the coherent spatial pattern observed in multiple independent genomic loci, with the two alternative haplogroups being differentially fixed in the Western Canadian Arctic and Sweden. As we expect genetic variation in these two regions to be respectively associated with the Arctic and Atlantic lineages, this local PCA approach thus reveals putative local ancestry tracts for either lineage in admixed individuals, which were found mostly in the Southern Canadian Arctic and Western Greenland. This constitutes a rather indirect method compared to phasing algorithms based on the Li and Stephens model [41] which requires a reference panel (e.g. HAPMIX [42] and FLARE [43]). However, no solution currently exists for the accurate phasing of low-coverage unlinked short-read sequencing data. By exploring how local PCAs highlight local genetic patterns coherent with haplotype structure, we hope to spur interest in improving how lcWGS can be used to their fullest.

Here, we only scratched the surface of the haplotypic structure and likely had the power to detect only the longest and most geographically structured haploblocks. A quick look at the structure outside MDS outlier regions (Fig. 5a, Fig S8) shows how most of the genome segregates similarly at a finer scale, i.e. in smaller blocks. The delimitation of haploblocks could likely be refined by tuning the window size and filtering parameters of the MDS outlier detection. However, some of the imprecision might be linked to reasons both technical (e.g., the uncertainty inherent to the analytical framework of low-coverage data) and biological (the heterogeneity of recombination along the genome and among populations, breaking ancestry tracks into regions of varying lengths). In the next section, we discuss one possible application for such haplotypic information in datasets of large sample sizes.

### Leveraging local ancestry to disentangle multiple origins of recolonization

In cases of post-glacial secondary contact, pinpointing the origin of recolonization – i.e. the position of the glacial refugium – of each lineage can help provide additional context, as the process of recolonization itself and its direction can have predictable impacts on within-lineage genetic variation [13,44]. One such impact is increased genetic drift through sequential founder effects along the recolonization front [45–47], leading to a decrease in genetic diversity and an accumulation of derived and sometimes deleterious alleles via increased genetic drift [48–50]. The directionality index (*ψ*) and TDoA approach [51] uses this prediction to triangulate the origin of recent range expansions by comparing the spatial distribution of derived alleles (e.g. [52–54]).

In our data, applying this method to the genome-wide variation pointed toward an origin in the south-eastern Canadian Arctic. However, the study system shows clear signs of admixture between lineages of at least two distinct origins, as is often the case in similar studies [55,56]. This violates a major assumption of the model [51], as the high divergence between lineages most likely overshadows any within-lineage signal of directionality. The Arctic lineage is expected to have recolonized from a smaller source population than other lineages [22], suggesting a demographic bottleneck that could explain the higher frequency of derived alleles in the northern, non-admixed populations. With the goal of revealing patterns specific to the Arctic lineage, we focused on genetic variation inside putative local ancestry tracts identified earlier. By considering only homozygotes for the haplogroup that we attributed to the Arctic lineage, we hypothesized that we could limit the influence of the introgression from the Atlantic lineage and investigate the origin of recolonization of the Arctic lineage. While these inferences had lower explanatory power and replicability, they globally pointed in a direction opposite to the genome-wide analysis, highlighting the interest in dissecting horizontal information present in genomes and further refinement of local ancestry tracts.

Inferring range expansions from the distribution of derived alleles has several limitations, as other evolutionary mechanisms can create asymmetry in two-dimensional site-frequency spectra, such as demographic changes unrelated to the expansion [57]. Boundary effects can also act as confounding factors, as populations on the outer part of the distribution will experience increased drift compared to central ones, which can create clines in *ψ* independently from range expansions [58]. In light of this and in the absence of rigorous hypothesis testing, our key takeaway focuses less on our confidence in the inference for this specific system and more on the potential of genetic information preserved in large blocks of known local ancestry to unravel complex, multi-layered evolutionary histories.

Despite previously discussed limitations, other sources of data were consistent with a post-glacial recolonization from the northwestern Canadian Arctic, such as mitochondrial diversity being higher in this region than elsewhere in the range of the Arctic lineage. While species are intuitively expected to have survived glacial periods in refugia south of the ice extent, fossil and genetic evidence suggests that higher-latitude refugia might have existed [14,56,59–61]. Although recent reconstructions suggest a complete glaciation of the Arctic archipelago at the last maximum (Fig. 1) [62], data from rodents [63], birds [64], and plants [65,66] point to cryptic refugia in the High Arctic, and Banks Island (Inuvialuit, North West Territories) has been proposed as a refugium for caribou [67]. Ultimately, the use of local ancestry as a lens through which to study recolonization could offer a promising avenue for disentangling complex post-glacial evolutionary histories of the High Arctic and elsewhere.

## Conclusion

Until recently, applications of whole-genome sequencing (WGS) to population genomics have been limited by high costs, resulting in low sample sizes and restricted study areas. Here, our use of low-coverage WGS data showcases how versatile this approach can be in investigating intraspecific evolutionary processes for widely sampled organisms. The inclusion of cytoplasmic DNA (from mitochondria or chloroplasts in plants) in WGS also promises a more systematic comparison of markers from different independently evolving segments of the genome, providing new phylogeographic insight while integrating earlier research in long-studied systems. Finally, WGS data offers far more than an increased SNP count, and we are confident that the scientific community will benefit from exploring innovative approaches for leveraging horizontal genetic signals, like linkage and recombination, to deepen our understanding of evolutionary dynamics.

## Material & methods

### Library preparation, sequencing, and data preprocessing

Anadromous Arctic Char adipose fins were collected at the mouth of rivers across Arctic Canada and Western Greenland between 2007 and 2023 and stored in ethanol 95% (4°C) or RNAlater (−20°C). Whole-genome libraries were prepared following Therkildsen & Palumbi (2017). Briefly, DNA was extracted using Nucleomag kits, cleaned with Axygen magnetic beads, quantified with AccuClear Ultra High Sensitivity dsDNA Quantification kits, and diluted to 1.5 ng/µl. Sample extracts were then used for Nextera library preparation: tagmentation (i.e. fragmentation and addition of adapters), 2 PCR steps to attach barcodes, and size selection for fragments 400-700bp in length using Axygen magnetic beads. To achieve a final depth of coverage of 2X, we performed sequencing on lanes of Illumina HiSeq 4000, with around 96 individual libraries per lane (range of 75 to 106), yielding on average 31.6 million reads per sample for 1178 samples in 33 populations. For the medium-coverage dataset, we selected 100 random samples in 5 populations (20 per population) and repeated sequencing on 5 lanes to obtain 135.9 million reads per sample (adding ∼6X).

All data were prepared following the pipeline described at https://github.com/enormandeau/wgs_sample_preparation. In brief, raw sequences were trimmed using fastp [68] and aligned on the ASM291031v2 reference genome with bwa mem (minimum alignment quality of 10) [69]. Note that the reference genome used was assembled using sequencing data from a Dolly Varden (*Salvelinus malma*) or a *S. alpinus* x *S. malma* hybrid [70] but remains the closest high-quality reference for Arctic Char at the time of our analysis. Duplicate reads were then removed using picard, indels were realigned, and overlapping ends of paired reads were clipped. The average per-base depth of coverage was then estimated using mosdepth [71], and 61 samples with genome-wide coverage under 1X were excluded from analyses on nuclear DNA. For the 5 populations targeted by additional sequencing, the 2X and 6X aligned reads were merged so that these populations could be analyzed at a coverage of either 2 or 8X.

### mtDNA and nuDNA genotyping and filtering

The depth of coverage was much higher on the mitochondrial genome, allowing us to call haplotypes for the whole mitogenome (16,659 bp) for every individual in the 2X dataset using angsd -doFasta 1 [72]. Sequences were cleaned in Geneious 2022.0.1 (https://www.geneious.com) by removing positions, then individuals, with more than 5% missing data. We used popart [73] to construct a median-joining network of haplotypes extracted for the D-loop control region (998 bp), using an ε parameter of zero. An individual sampled in population 30 matched the D-loop sequence for *Salvelinus fontinalis* and was kept as an outgroup for the haplotype network but was removed from every subsequent analysis.

Nuclear SNP positions were first detected using ANGSD v0.937 [72] (-GL 2 - remove_bads 1 -minMapQ 30 -minQ 20 -skipTriallelic 1 -uniqueOnly 1 -only_proper_pairs 1 -min_Maf = 0.01) on both the 2X and 8X datasets independently with the following coverage filters (for 2X data: at least 1X in 75% of samples, maximum average depth of 4X; for 8X data: at least 3X in 75% of samples, maximum average depth of 20X). The sex of sampled fish was unknown. As such, SNPs on sex-linked regions identified by Beemelmanns et al. (2024) were not considered, to avoid potentially unbalanced sex ratios being confounded with population differentiation.

The 15.6 (2X) and 21.5 (8X) million retained SNP positions were tested for their deviation of Mendelian inheritance using the calcLR in ngsParalog [74]. ngsParalog likelihood ratios were compared to a **χ**^2^ distribution and SNPs with a P-value (adjusted by applying the Bonferroni procedure) under the threshold of 0.001 were considered as deviant (2X: 9.63 million, 62% of all SNPs; 8X: 11.20 million, 52%). Deviant SNPs were removed from the site list for each dataset, as they are expected to have their inferred genotype biased by mismapped reads related to paralogy or repeated elements [75]. For analyses based on site frequency spectra (SFS), we masked the reference genome in 150bp regions centered on every deviant SNP, as well as all coding genes and a 1-kb region around them.

We estimated genotype likelihoods for 5.98 million non-deviant SNPs in the 2X dataset using -doGlf -GL2 in ANGSD. We used ngsLD to estimate linkage between SNPs within 500 kb using a random subset of half the samples and pruned the dataset with the graph method until no SNP pairs within 200kb had an r^2^ above 0.1, leaving 375,236 SNPs in the LD-pruned dataset.

### Population structure, differentiation, and diversity

The general distribution of genetic variation and population structure was first assessed by performing a principal components analysis on the LD-pruned 2X dataset in PCAngsd 1.10 [76]. We then explored the hierarchical population structure using NGSadmix [77] on the LD-pruned 2X data to estimate individual admixture while varying the number of ancestral populations (K) from 2 to 33.

Pairwise population differentiation was estimated using allele frequency difference (AFD) and F_ST_ index. For AFD, we estimated allele frequency in every population for all 5.98 million SNPs (2X) using angsd -domaf 1, then calculated the average absolute MAF difference between each pair of populations. For F_ST_, we used ANGSD and the reference genome masked for deviant SNPs and genes to estimate sample allele frequency (SAF) for each population, then we used the winSFS streaming option [78] to compute two-population SFS while accounting for linkage disequilibrium. The SAF and 2D-SFS served as input for the *realsfs fst index*, *print*, *stats,* and *stats2* functions in ANGSD to calculate genome-wide weighted F_ST_ [79]. We tested for the presence of isolation-by-distance through linear regression between the marine distance between population pairs, estimated using marmap [80], and either their AFD or linearized F_ST_ [81].

We estimated nuclear genetic diversity in populations with Watterson’s estimator (ϴ_W_) and nucleotide diversity (ϴ_π_), as well as individual heterozygosity. ϴ statistics were computed with ANGSD’s *realsfs saf2theta* and *thetaStat do_stat* using the SAF by population described above and a 1D-SFS produced by winsfs. To account for the variable coverage along the genome and across populations, we compared per-site ϴ in windows of 100 kb (step of 10 kb) and excluded windows with a number of sites (denominator) under the 5th percentile (2,600 sites) in any population. Individual heterozygosity was estimated by computing a per-sample SAF and dividing the number of heterozygous sites (SAF = 1) by the number of sites.

We also estimated diversity metrics on whole mitogenome sequences, after removing haplotypes from the Atlantic lineage, those with more than 5% of missing data, and masking positions with remaining missing values. The 1,074 Arctic haplotypes were 16,483 bp long and were analyzed by population with the *pegas* [82] package in R for 1) ϴ_W_ with *theta.s* function, 2) ϴ_π_ with the *nuc.div* function, 3) Tajima’s D with the Tajima.test function, 4) haplotype diversity using the *haploFreq* function and the formula corrected for sample size from [83], and 5) the frequency of the Arctic haplotype most frequent in the whole dataset. We tested for geographical patterns in populational mtDNA diversity using regression models for those 5 variables with the marine distance from the westernmost population (pop.1) as the explicative variable. We used the Akaike Information Criterion (AIC) [84] to compare the null, simple, and 2nd-degree polynomial models.

### Secondary contact, introgression, and demographic inference

To test hypotheses regarding the admixture between the Arctic and Atlantic lineage of Arctic Char in Canada, we downloaded *S. alpinus* raw sequences (PRJEB62707) from an inland lake in Sweden (Lilla Stensjön) [85], which we assumed represents a population descending purely from the Atlantic lineage. We also acquired unpublished data from a Dolly Varden (*Salvelinus malma*) population in Babage River, Yukon, Canada (Labrecque et al., in prep). Those sequences were trimmed, aligned, and cleaned as described in the “data preprocessing” section, then we estimated genotype likelihoods and population MAF for the LD-pruned SNP list. We obtained an average depth of coverage of 5X for the Swedish population and 2X for *S. malma*.

We repeated the NGSadmix analysis for K=2 with the Swedish population. We converted the estimated MAF of all 35 populations of *S. alpinus* and *S. malma* to approximate allele count by multiplying the MAF by the number of sampled alleles (twice the sample size) to fit the Treemix file format. We computed f_4_ statistics in Treemix 1.13 [86] with *S. malma* as the outgroup (H_4_), the Swedish population as the potential introgressor (H_3_), and all pairs of Canadian and Greenlandic populations as H_1_ and H_2_.

Based on the previous results, we build alternative demographic models to test the presence of ancient or recent gene flow between the Atlantic lineage, represented by the Swedish population and the Southern Arctic Canada (see Fig. 4B). We estimated the unfolded 3D-SFS between COP (North, pop. 4), TJL (South, pop. 25), and LLS (Sweden, pop. 34) using winsfs. Alleles were polarized based on an ancestral reference genome constructed by aligning WGS data from four closely related species (Atlantic Salmon; Lake Trout, *Salvelinus namaycush*; Rainbow Trout, *Oncorhynchus mykiss*; and Chinook Salmon, *Oncorhynchus tshawytscha*) on the *Salvelinus sp.* reference genome (ASM291031v2), as in [75]. We used the most common allele to create a consensus for each species using *angsd -doFasta*, then merged the consensus in Geneious 2022.0.1 to keep only positions covered in at least 2 consensuses. The fit between the observed SFS and models with 0) no, 1) ancient, and 2) recent gene flow was compared by running 100 runs of each in fastsimcoal 2.7 with the following parameters: -d -C 10 -n 500000 - L 50 -s 0 -M. [Parameter estimation]

### Introgression and recombination landscapes

We identified genomic regions displaying distinct genetic variation from the genome-wide signal using local PCAs as described in [38,40]. Briefly, we scanned each chromosome independently by computing the covariance matrix between individuals in non-overlapping windows of 100 SNPs in PCAngsd and detecting clusters of consecutive windows presenting similar PCA patterns using the *pcdist* function and multidimensional scaling (MDS) in the R package *lostruct*. Windows with MDS values beyond 3 standard deviations from the chromosomal mean on one of the first five MDS axes were kept, and clusters within 20 windows of each other were merged. We only considered clusters longer than 10 windows.

In each outlier cluster, we tested for the presence of patterns on PC1 indicative of the three genotypes of a chromosomal inversion or other types of nonrecombinant haploblock. The R package *dbscan* allowed for the grouping of individuals based on their density distribution on PC1 while allowing for uncategorized individuals between clusters. We iteratively ran *dbscan* while increasing the ε parameter by increments of 0.005, starting from 0.01, until three groups were detected or ε = 0.15 was reached. We further cleaned the *dbscan* results by setting individuals with PC1 values outside 3 standard deviations from the mean of any group as intermediate genotype. The three remaining groups were labeled haplogroup AA, AB, and BB, starting from the most frequent one, and the intermediate genotypes were labelled A0 and 0B.

To test if the putative haploblocks were associated to lower recombination rates, we estimated recombination using LDhat for each chromosome and population in the 8X dataset. To produce the bcf file in input, we genotyped previously identified non-deviant SNPs in ANGSD using -doBcf 1 -doGeno -4 -doPost 1 -postCutoff 0.95 with more stringent filters (population MAF > 0.05, minimum of 4X in 90% of individuals). Population-scaled recombination rates (*ρ*/kb) was estimated in LDhat 2.2 [87] while controlling for demography with LDpop [88] using a θ estimate of 0.001, a block penalty of 5 and other parameters set following the pipeline in [89]. The LDhat output was cleaned by removing plateaus of invariable recombining rates longer than 22 kb (99^th^ percentile of the length of all plateaus). The weighted mean of recombination rates inside each putative haploblocks and other MDS outliers was compared to the chromosome-wide average through mixed-effect models with the chromosome and population set as random effects. Chromosomes and haploblocks were only considered in recombination maps that covered at least 60% of their length.

### Inference of the origin of recolonization

To triangulate the origin of the recolonization after the last glacial maximum, and thus a putative glacial refugium, we computed the directionality index (ψ) for each population pair from their genome-wide unfolded 2D-SFS, following Peter & Slatkin (2013). Ancestral alleles private to either population were excluded. We then used the Time Difference of Arrival algorithm [90] on the ψ matrix following Prior et al. (2020). Briefly, a pairwise comparison of ψ and geographical distance between focal populations and the assumed origin was performed for every possible origin, using a 0.5 decimal degrees step across the study area. For origins with a positive slope between ψ and geographic distance, we rescaled the coefficient of regression (R^2^) from 0 to 1, and origins with a higher R^2^ were considered more likely. For this analysis, we excluded populations out of Canada, as well as population 30, which had a high level of Atlantic lineage ancestry.

To test how introgression from the Atlantic lineage impacted the inference of the origin of recolonization, we repeated the *ψ* and TDoA analyses while focusing on the diversity inside the Arctic alleles for the putative haploblocks identified using local PCAs. For each population pair and haploblock, we randomly selected 10 homozygotes for Arctic allele (AA) by population and computed their SAFs and 2D-SFS. We then summed the per-haploblock SFS where both populations had at least 10 AA individuals and computed a directionality index from the summed spectrum. TDoA was performed on the resulting ψ matrix as described above.

To assess the robustness of this analysis, we repeated this process over 100 replicates on resampled AA individuals and a random subset of 90% of putative haploblocks, as well as over 100 replicates where each haploblock was replaced by a random genomic region of equal length on the same chromosome. TDoA R^2^ maps from the putative haploblocks and random regions were each rescaled as above, then averaged and compared to the genome-wide TDoA signal.

## Data Accessibility Statement

Part of the *Salvelinus alpinus* sequencing data analyzed in this manuscript is available on Short Read Archive as part of projects PRJNA1031558 (Canada and Greenalnd) and PRJEB62707 (Sweden). Following submission, all new genetic data will be deposited on SRA. Sequencing data on *Salvelinus malma* is also planned to be deposited as part of a separate publication, presently in preparation.

## Acknowledgment

This work was supported by a Large-Scale Applied Research Project grant from Genome Canada named “FISHES: Fostering Small-scale Fisheries for Health, Economy and Food Security”. Sampling was made possible by the collaboration of Fisheries and Ocean Canada; Ministère de l’Environnement, de la Lutte contre les changements climatiques, de la Faune et des Parcs (Québec; Julien Mainguy); Government of Nunavut; Makivik Corporation (Nunavik); Aarhus University (Denmark; Magnus W. Jacobsen); and numerous Inuit communities, organizations, and local fishers across Canada and Greenland. Many thanks to Anne Beemelmanns, Bérénice Bougas, Charles Babin, Isabeau Caza-Allard, Louis-Philippe Collin, Alysse Perreault-Payette, and Gabriel Piette-Lauzière for their help with laboratory work and coordination, as well as Raphaël Bouchard, Nicolas Bierne, Lila Colston-Nepali, Pierre-Alexandre Gagnaire, Laura Meyer, Adrien Tran Lu Y, Quentin Rougemont, and Florent Sylvestre for their support and suggestions during the investigation of the data. We also thank Rennes Metropole, ECOBIO (Université de Rennes), and ISEM (Université de Montpellier) for providing funding or access to their facility which were invaluable in the completion of this work.

## References

1. Avise JC, Arnold J, Ball RM, Bermingham E, Lamb T, Neigel JE, et al. Intraspecific Phylogeography: The Mitochondrial DNA Bridge Between Population Genetics and Systematics. Annu Rev Ecol Syst. 1987;18:489–522.

2. Edwards SV, Shultz AJ, Campbell-Staton SC. Next-generation sequencing and the expanding domain of phylogeography. Folia Zool [Internet]. 2015 Nov [cited 2024 Jun 27];64(3):187–206. Available from: http://www.bioone.org/doi/10.25225/fozo.v64.i3.a2.2015

3. Ballard JWO, Whitlock MC. The incomplete natural history of mitochondria. Mol Ecol [Internet]. 2004 [cited 2024 Nov 25];13(4):729–44. Available from: https://onlinelibrary.wiley.com/doi/abs/10.1046/j.1365-294X.2003.02063.x

4. Toews DPL, Brelsford A. The biogeography of mitochondrial and nuclear discordance in animals. Mol Ecol [Internet]. 2012 Aug [cited 2024 Apr 23];21(16):3907–30. Available from: https://onlinelibrary.wiley.com/doi/10.1111/j.1365-294X.2012.05664.x

5. Zink RM, Barrowclough GF. Mitochondrial DNA under siege in avian phylogeography. Mol Ecol [Internet]. 2008 May [cited 2024 Jun 27];17(9):2107–21. Available from: https://onlinelibrary.wiley.com/doi/10.1111/j.1365-294X.2008.03737.x

6. Peter BM. A geometric relationship of *F* _2_ , *F* _3_ and *F* _4_ -statistics with principal component analysis. Philos Trans R Soc B Biol Sci [Internet]. 2022 Jun 6 [cited 2024 Aug 23];377(1852):20200413. Available from: https://royalsocietypublishing.org/doi/10.1098/rstb.2020.0413

7. Bhatia G, Patterson N, Sankararaman S, Price AL. Estimating and interpreting *F* _ST_ : The impact of rare variants. Genome Res [Internet]. 2013 Sep [cited 2024 Aug 23];23(9):1514–21. Available from: http://genome.cshlp.org/lookup/doi/10.1101/gr.154831.113

8. Leitwein M, Duranton M, Rougemont Q, Gagnaire PA, Bernatchez L. Using Haplotype Information for Conservation Genomics. Trends Ecol Evol [Internet]. 2020 Mar [cited 2024 Mar 29];35(3):245–58. Available from: https://linkinghub.elsevier.com/retrieve/pii/S0169534719303040

9. Martin AR, Gignoux CR, Walters RK, Wojcik GL, Neale BM, Gravel S, et al. Human Demographic History Impacts Genetic Risk Prediction across Diverse Populations. Am J Hum Genet [Internet]. 2017 Apr 6 [cited 2024 Aug 28];100(4):635–49. Available from: https://www.cell.com/ajhg/abstract/S0002-9297(17)30107-6

10. Shipilina D, Pal A, Stankowski S, Chan YF, Barton NH. On the origin and structure of haplotype blocks. Mol Ecol [Internet]. 2023 Mar [cited 2024 Jun 25];32(6):1441–57. Available from: https://onlinelibrary.wiley.com/doi/10.1111/mec.16793

11. Lawson DJ, Hellenthal G, Myers S, Falush D. Inference of Population Structure using Dense Haplotype Data. PLOS Genet [Internet]. 2012 Jan 26 [cited 2024 Oct 11];8(1):e1002453. Available from: https://journals.plos.org/plosgenetics/article?id=10.1371/journal.pgen.1002453

12. Hewitt GM. Some genetic consequences of ice ages, and thier role in divergence. Biol J Linn Soc. 1996;58(September 1994):247–76.

13. Hewitt GM. The genetic legacy of the quaternary ice ages. Nature. 2000;405(6789):907–13.

14. Shafer ABA, Cullingham CI, Côté SD, Coltman DW. Of glaciers and refugia: a decade of study sheds new light on the phylogeography of northwestern North America. Mol Ecol. 2010 Nov;19(21):4589–621.

15. Bernatchez L, Wilson CC. Comparative phylogeography of Nearctic and Palearctic fishes. Mol Ecol. 1998;7(4):431–52.

16. Hewitt GM. Post-glacial re-colonization of European biota. Biol J Linn Soc [Internet]. 1999 Sep 1 [cited 2024 Aug 23];68(1–2):87–112. Available from: 10.1111/j.1095-8312.1999.tb01160.x

17. Liang M, Nielsen R. The Lengths of Admixture Tracts. Genetics [Internet]. 2014 Jul 1 [cited 2024 Aug 13];197(3):953–67. Available from: 10.1534/genetics.114.162362

18. Wiens BJ, Colella JP. That’s not a Hybrid: How to Distinguish Patterns of Admixture and Isolation-by-Distance [Internet]. 2024 [cited 2024 Apr 23]. Available from: http://biorxiv.org/lookup/doi/10.1101/2024.04.15.589658

19. Hocutt CH, Wiley EO. The Zoogeography of North American freshwater fishes [Internet]. New York: Wiley; 1986 [cited 2024 Nov 25]. 866 p. Available from: https://bac-lac.on.worldcat.org/oclc/299840901

20. Klemetsen A. The Charr Problem Revisited: Exceptional Phenotypic Plasticity Promotes Ecological Speciation in Postglacial Lakes. Freshw Rev. 2010;3(1):49–74.

21. Brunner PC, Douglas MR, Osinov A, Wilson CC, Bernatchez L. Holarctic Phylogeography of Arctic Charr (Salvelinus Alpinus L.) Inferred From Mitochondrial Dna Sequences. Evolution [Internet]. 2001;55(3):573–86. Available from: http://doi.wiley.com/10.1111/j.0014-3820.2001.tb00790.x

22. Moore JS, Bajno R, Reist JD, Taylor EB. Post-glacial recolonization of the North American Arctic by Arctic char (Salvelinus alpinus): Genetic evidence of multiple northern refugia and hybridization between glacial lineages. J Biogeogr. 2015;42(11):2089–100.

23. Dallaire X, Normandeau É, Mainguy J, Tremblay JÉ, Bernatchez L, Moore JS. Genomic data support management of anadromous Arctic Char fisheries in Nunavik by highlighting neutral and putatively adaptive genetic variation. Evol Appl [Internet]. 2021 [cited 2021 Jun 7];n/a(n/a). Available from: https://onlinelibrary.wiley.com/doi/abs/10.1111/eva.13248

24. Gordeeva NV, Alekseyev SS, Kirillov AF, Romanov VI, Pichugin MYu. New Data about the Distribution of Three Phylogenetic Lineages of Arctic Charr Salvelinus alpinus (Salmonidae) in their Contact Zones in the North of East Siberia. J Ichthyol [Internet]. 2021 Sep [cited 2024 Apr 23];61(5):701–8. Available from: https://link.springer.com/10.1134/S0032945221050064

25. Jacobsen MW, Jensen NW, Nygaard R, Præbel K, Jónsson B, Nielsen NH, et al. A melting pot in the Arctic: Analysis of mitogenome variation in Arctic char (*Salvelinus alpinus*) reveals a 1000-km contact zone between highly divergent lineages. Ecol Freshw Fish [Internet]. 2022 Apr [cited 2023 Apr 3];31(2):330–46. Available from: https://onlinelibrary.wiley.com/doi/10.1111/eff.12633

26. Salisbury SJ, McCracken GR, Keefe D, Perry R, Ruzzante DE. Extensive secondary contact among three glacial lineages of Arctic Char (Salvelinus alpinus) in Labrador and Newfoundland. Ecol Evol [Internet]. 2019;9(4):2031–45. Available from: http://doi.wiley.com/10.1002/ece3.4893

27. Moore JS, Harris LN, Tallman RF, B TE. The interplay between dispersal and gene flow in anadromous Arctic char (Salvelinus alpinus): implications for potential for local adaptation. Can J Fish Aquat Sci [Internet]. 2013;1338(July):1327–38. Available from: http://www.nrcresearchpress.com/doi/abs/10.1139/cjfas-2013-0138

28. Patterson N, Moorjani P, Luo Y, Mallick S, Rohland N, Zhan Y, et al. Ancient Admixture in Human History. Genetics [Internet]. 2012 Nov 1 [cited 2024 Oct 24];192(3):1065–93. Available from: 10.1534/genetics.112.145037

29. Reich D, Thangaraj K, Patterson N, Price AL, Singh L. Reconstructing Indian population history. Nature [Internet]. 2009 Sep [cited 2024 Oct 24];461(7263):489–94. Available from: https://www.nature.com/articles/nature08365

30. Galtier N, Nabholz B, Glémin S, Hurst GDD. Mitochondrial DNA as a marker of molecular diversity: a reappraisal. Mol Ecol [Internet]. 2009 [cited 2024 Aug 28];18(22):4541–50. Available from: https://onlinelibrary.wiley.com/doi/abs/10.1111/j.1365-294X.2009.04380.x

31. Hudson RR, Turelli M. STOCHASTICITY OVERRULES THE “THREE-TIMES RULE”: GENETIC DRIFT, GENETIC DRAFT, AND COALESCENCE TIMES FOR NUCLEAR LOCI VERSUS MITOCHONDRIAL DNA. Evolution [Internet]. 2003 Jan [cited 2024 Aug 26];57(1):182–90. Available from: https://academic.oup.com/evolut/article/57/1/182/6755810

32. Dempson JB, Kristofferson AH. Spatial and Temporal Aspects of the Ocean Migration of Anadromous Arctic Char. Am Fish Soc Symp. 1987;1:340–57.

33. Moore JS, Harris LN, Kessel ST, Bernatchez L, Tallman RF, Fisk AT. Preference for near-shore and estuarine habitats in anadromous Arctic char (Salvelinus alpinus) from the Canadian high Arctic (Victoria Island, NU) revealed by acoustic telemetry. Can J Fish Aquat Sci. 2016;53(9):1689–99.

34. Lou RN, Jacobs A, Wilder A, Therkildsen NO. A beginner’s guide to low-coverage whole genome sequencing for population genomics [Internet]. Preprints; 2021 Apr [cited 2021 May 6]. Available from: https://www.authorea.com/users/380682/articles/496574-a-beginner-s-guide-to-low-coverage-whole-genome-sequencing-for-population-genomics?commit=a25912ea40ed4aa7f84aeab0b40f1e4d2bea282c

35. Rubinacci S, Ribeiro DM, Hofmeister RJ, Delaneau O. Efficient phasing and imputation of low-coverage sequencing data using large reference panels. Nat Genet [Internet]. 2021 Jan [cited 2024 Aug 8];53(1):120–6. Available from: https://www.nature.com/articles/s41588-020-00756-0

36. Meier JI, Salazar PA, Kučka M, Davies RW, Dréau A, Aldás I, et al. Haplotype tagging reveals parallel formation of hybrid races in two butterfly species. Proc Natl Acad Sci [Internet]. 2021 Jun 22 [cited 2024 Nov 11];118(25):e2015005118. Available from: https://www.pnas.org/doi/full/10.1073/pnas.2015005118

37. Ishigohoka J, Bascón-Cardozo K, Bours A, Fuß J, Rhie A, Mountcastle J, et al. Distinct patterns of genetic variation at low-recombining genomic regions represent haplotype structure [Internet]. 2024 [cited 2024 Jun 21]. Available from: http://biorxiv.org/lookup/doi/10.1101/2021.12.22.473882

38. Huang K, Andrew RL, Owens GL, Ostevik KL, Rieseberg LH. Multiple chromosomal inversions contribute to adaptive divergence of a dune sunflower ecotype. Mol Ecol [Internet]. 2020 [cited 2021 Aug 9];29(14):2535–49. Available from: https://onlinelibrary.wiley.com/doi/abs/10.1111/mec.15428

39. Hale MC, Campbell MA, McKinney GJ. A candidate chromosome inversion in Arctic charr (Salvelinus alpinus) identified by population genetic analysis techniques. Macqueen D, editor. G3 GenesGenomesGenetics [Internet]. 2021 Jul 28 [cited 2021 Aug 30];jkab267. Available from: https://academic.oup.com/g3journal/advance-article/doi/10.1093/g3journal/jkab267/6329827

40. Mérot C, Berdan EL, Cayuela H, Djambazian H, Ferchaud AL, Laporte M, et al. Locally Adaptive Inversions Modulate Genetic Variation at Different Geographic Scales in a Seaweed Fly. Charlesworth D, editor. Mol Biol Evol [Internet]. 2021 Aug 23 [cited 2024 Sep 5];38(9):3953–71. Available from: https://academic.oup.com/mbe/article/38/9/3953/6272233

41. Li N, Stephens M. Modeling linkage disequilibrium and identifying recombination hotspots using single-nucleotide polymorphism data. Genetics [Internet]. 2003 Dec [cited 2024 Dec 4];165(4):2213–33. Available from: https://www.ncbi.nlm.nih.gov/pmc/articles/PMC1462870/

42. Price AL, Tandon A, Patterson N, Barnes KC, Rafaels N, Ruczinski I, et al. Sensitive Detection of Chromosomal Segments of Distinct Ancestry in Admixed Populations. PLOS Genet [Internet]. 2009 Jun 19 [cited 2024 Aug 28];5(6):e1000519. Available from: https://journals.plos.org/plosgenetics/article?id=10.1371/journal.pgen.1000519

43. Browning SR, Waples RK, Browning BL. Fast, accurate local ancestry inference with FLARE. Am J Hum Genet [Internet]. 2023 Feb 2 [cited 2024 Aug 28];110(2):326–35. Available from: https://www.ncbi.nlm.nih.gov/pmc/articles/PMC9943733/

44. Peischl S, Kirkpatrick M, Excoffier L. Expansion Load and the Evolutionary Dynamics of a Species Range. Am Nat [Internet]. 2015 Apr [cited 2021 Jun 3];185(4):E81–93. Available from: https://www.journals.uchicago.edu/doi/10.1086/680220

45. Braga RT, Rodrigues JFM, Diniz-Filho J a. F, Rangel TF. Genetic Population Structure and Allele Surfing During Range Expansion in Dynamic Habitats. An Acad Bras Ciênc [Internet]. 2019 Apr 25 [cited 2024 Oct 8];91:e20180179. Available from: https://www.scielo.br/j/aabc/a/bjgBZQhBLJh5MFQNfC8v9Qh/?lang=en

46. Hallatschek O, Hersen P, Ramanathan S, Nelson DR. Genetic drift at expanding frontiers promotes gene segregation. Proc Natl Acad Sci U S A. 2007;104(50):19926–30.

47. Slatkin M, Excoffier L. Serial Founder Effects During Range Expansion: A Spatial Analog of Genetic Drift. Genetics [Internet]. 2012 May [cited 2021 Jun 2];191(1):171–81. Available from: https://www.ncbi.nlm.nih.gov/pmc/articles/PMC3338258/

48. Koski MH, Layman NC, Prior CJ, Busch JW, Galloway LF. Selfing ability and drift load evolve with range expansion. Evol Lett [Internet]. 2019 [cited 2021 Jun 2];3(5):500–12. Available from: https://onlinelibrary.wiley.com/doi/abs/10.1002/evl3.136

49. Rougemont Q, Moore JS, Leroy T, Normandeau E, Rondeau EB, Withler RE, et al. The role of historical contingency in shaping geographic pattern of deleterious mutation load in a broadly distributed Pacific Salmon. 2020; Available from: http://www.xkcd.com/446/#

50. Rougemont Q, Leroy T, Rondeau EB, Koop B, Bernatchez L. Allele surfing causes maladaptation in a Pacific salmon of conservation concern. PLOS Genet [Internet]. 2023 Sep 8 [cited 2024 Oct 8];19(9):e1010918. Available from: https://journals.plos.org/plosgenetics/article?id=10.1371/journal.pgen.1010918

51. Peter BM, Slatkin M. Detecting Range Expansions from Genetic Data. Evolution [Internet]. 2013 [cited 2021 Jun 8];67(11):3274–89. Available from: https://onlinelibrary.wiley.com/doi/abs/10.1111/evo.12202

52. Ioannidis AG, Blanco-Portillo J, Sandoval K, Hagelberg E, Barberena-Jonas C, Hill AVS, et al. Paths and timings of the peopling of Polynesia inferred from genomic networks. Nature [Internet]. 2021 Sep [cited 2024 Oct 11];597(7877):522–6. Available from: https://www.nature.com/articles/s41586-021-03902-8

53. Luqman H, Wegmann D, Fior S, Widmer A. Climate-induced range shifts drive adaptive response via spatio-temporal sieving of alleles. Nat Commun [Internet]. 2023 Feb 25 [cited 2024 Oct 11];14(1):1080. Available from: https://www.nature.com/articles/s41467-023-36631-9

54. Prior CJ, Layman NC, Koski MH, Galloway LF, Busch JW. Westward range expansion from middle latitudes explains the Mississippi River discontinuity in a forest herb of eastern North America. Mol Ecol [Internet]. 2020 Nov [cited 2024 May 6];29(22):4473–86. Available from: https://onlinelibrary.wiley.com/doi/10.1111/mec.15650

55. Nugent CM, Kess T, Langille BL, Beck SV, Duffy S, Messmer A, et al. Post-glacial recolonization and multiple scales of secondary contact contribute to contemporary Atlantic salmon (Salmo salar) genomic variation in North America. J Biogeogr [Internet]. 2024 [cited 2024 Oct 15];51(9):1767–82. Available from: https://onlinelibrary.wiley.com/doi/abs/10.1111/jbi.14852

56. Sommer RS, Zachos FE. Fossil evidence and phylogeography of temperate species: ‘glacial refugia’ and post-glacial recolonization. J Biogeogr [Internet]. 2009 [cited 2024 Sep 4];36(11):2013–20. Available from: https://onlinelibrary.wiley.com/doi/abs/10.1111/j.1365-2699.2009.02187.x

57. Mestre F, Barbosa S, Garrido-García JA, Pita R, Mira A, Alves PC, et al. Inferring past refugia and range dynamics through the integration of fossil, niche modelling and genomic data. J Biogeogr [Internet]. 2022 [cited 2024 Sep 4];49(11):2064–76. Available from: https://onlinelibrary.wiley.com/doi/abs/10.1111/jbi.14492

58. Kemppainen P, Schembri R, Momigliano P. Boundary Effects Cause False Signals of Range Expansions in Population Genomic Data. Mol Biol Evol [Internet]. 2024 May 1 [cited 2024 Sep 4];41(5):msae091. Available from: 10.1093/molbev/msae091

59. Mee JA, Moore JS. The ecological and evolutionary implications of microrefugia. J Biogeogr [Internet]. 2014 [cited 2024 Oct 24];41(5):837–41. Available from: https://onlinelibrary.wiley.com/doi/abs/10.1111/jbi.12254

60. Premoli A, Mathiasen P, Kitzberger T. Southern-most Nothofagus trees enduring ice ages: Genetic evidence and ecological niche retrodiction reveal high latitude (54°S) glacial refugia. Palaeogeogr Palaeoclimatol Palaeoecol. 2010 Dec 1;298:247–56.

61. Wójcik JM, Kawałko A, Marková S, Searle JB, Kotlík P. Phylogeographic signatures of northward post-glacial colonization from high-latitude refugia: a case study of bank voles using museum specimens. J Zool [Internet]. 2010 [cited 2024 Sep 4];281(4):249–62. Available from: https://onlinelibrary.wiley.com/doi/abs/10.1111/j.1469-7998.2010.00699.x

62. Dalton AS, Margold M, Stokes CR, Tarasov L, Dyke AS, Adams RS, et al. An updated radiocarbon-based ice margin chronology for the last deglaciation of the North American Ice Sheet Complex. Quat Sci Rev. 2020;234.

63. Fedorov, Stenseth. Multiple glacial refugia in the North American Arctic: inference from phylogeography of the collared lemming (Dicrostonyx groenlandicus) [Internet]. 2002 [cited 2024 Sep 4]. Available from: https://royalsocietypublishing-org.acces.bibl.ulaval.ca/doi/epdf/10.1098/rspb.2002.2126

64. Holder K, Montgomerie R, Friesen VL. A TEST OF THE GLACIAL REFUGIUM HYPOTHESIS USING PATTERNS OF MITOCHONDRIAL AND NUCLEAR DNA SEQUENCE VARIATION IN ROCK PTARMIGAN (LAGOPUS MUTUS). Evol Int J Org Evol. 1999 Dec;53(6):1936–50.

65. Abbott RJ, Smith LC, Milne RI, Crawford RMM, Wolff K, Balfour J. Molecular Analysis of Plant Migration and Refugia in the Arctic. Science [Internet]. 2000 [cited 2024 Sep 4];289(5483):1343–6. Available from: https://www.jstor.org/stable/3077629

66. Tremblay NO, Schoen DJ. Molecular phylogeography of Dryas integrifolia: glacial refugia and postglacial recolonization. Mol Ecol [Internet]. 1999 Jul [cited 2021 May 27];8(7):1187–98. Available from: http://doi.wiley.com/10.1046/j.1365-294x.1999.00680.x

67. Klütsch CFC, Manseau M, Anderson M, Sinkins P, Wilson PJ. Evolutionary reconstruction supports the presence of a Pleistocene Arctic refugium for a large mammal species. J Biogeogr. 2017;44(12):2729–39.

68. Chen S, Zhou Y, Chen Y, Gu J. fastp: an ultra-fast all-in-one FASTQ preprocessor. Bioinformatics [Internet]. 2018 Sep 1 [cited 2023 Jun 28];34(17):i884–90. Available from: https://academic.oup.com/bioinformatics/article/34/17/i884/5093234

69. Li H. Aligning sequence reads, clone sequences and assembly contigs with BWA-MEM [Internet]. arXiv.org. 2013 [cited 2024 Oct 3]. Available from: https://arxiv.org/abs/1303.3997v2

70. Christensen KA, Rondeau EB, Minkley DR, Leong JS, Nugent CM, Danzmann RG, et al. Retraction: The Arctic charr (Salvelinus alpinus) genome and transcriptome assembly. PLOS ONE [Internet]. 2021 Feb 9 [cited 2023 Oct 31];16(2):e0247083. Available from: https://journals.plos.org/plosone/article?id=10.1371/journal.pone.0247083

71. Pedersen BS, Quinlan AR. Mosdepth: quick coverage calculation for genomes and exomes. Hancock J, editor. Bioinformatics [Internet]. 2018 Mar 1 [cited 2024 Apr 24];34(5):867–8. Available from: https://academic.oup.com/bioinformatics/article/34/5/867/4583630

72. Korneliussen TS, Albrechtsen A, Nielsen R. ANGSD: Analysis of Next Generation Sequencing Data. BMC Bioinformatics [Internet]. 2014 Dec [cited 2021 May 7];15(1):356. Available from: https://bmcbioinformatics.biomedcentral.com/articles/10.1186/s12859-014-0356-4

73. Leigh JW, Bryant D. popart: full-feature software for haplotype network construction. Methods Ecol Evol [Internet]. 2015 [cited 2024 Dec 4];6(9):1110–6. Available from: https://onlinelibrary.wiley.com/doi/abs/10.1111/2041-210X.12410

74. Linderoth T. Identifying Population Histories, Adaptive Genes, and Genetic Duplication from Population-Scale Next Generation Sequencing. University of California, Berkeley; 2018.

75. Dallaire X, Bouchard R, Hénault P, Ulmo-Diaz G, Normandeau E, Mérot C, et al. Widespread Deviant Patterns of Heterozygosity in Whole-Genome Sequencing Due to Autopolyploidy, Repeated Elements, and Duplication. Genome Biol Evol [Internet]. 2023 Dec 1 [cited 2024 Oct 24];15(12):evad229. Available from: 10.1093/gbe/evad229

76. Meisner J, Albrechtsen A. Inferring Population Structure and Admixture Proportions in Low-Depth NGS Data. Genetics [Internet]. 2018 Oct 1 [cited 2024 Oct 24];210(2):719–31. Available from: 10.1534/genetics.118.301336

77. Skotte L, Korneliussen TS, Albrechtsen A. Estimating Individual Admixture Proportions from Next Generation Sequencing Data. Genetics [Internet]. 2013 Nov 1 [cited 2024 Apr 24];195(3):693–702. Available from: https://academic.oup.com/genetics/article/195/3/693/5935455

78. Rasmussen MS, Garcia-Erill G, Korneliussen TS, Wiuf C, Albrechtsen A. Estimation of site frequency spectra from low-coverage sequencing data using stochastic EM reduces overfitting, runtime, and memory usage. Gravel S, editor. Genetics [Internet]. 2022 Nov 30 [cited 2024 Apr 24];222(4):iyac148. Available from: https://academic.oup.com/genetics/article/doi/10.1093/genetics/iyac148/6730749

79. Reynolds J, Weir BS, Cockerham CC. Estimation of the Coancestry Coefficient: Basis for a Short-Term Genetic Distance. Genetics [Internet]. 1983 Nov 1 [cited 2024 May 7];105(3):767–79. Available from: https://academic.oup.com/genetics/article/105/3/767/5996242

80. Pante E, Simon-Bouhet B. marmap: A Package for Importing, Plotting and Analyzing Bathymetric and Topographic Data in R. PLoS ONE. 2013;8(9):6–9.

81. Slatkin M. Isolation by Distance in Equilibrium and Non-Equilibrium Populations. Evolution [Internet]. 1993 Feb [cited 2024 May 7];47(1):264–79. Available from: https://academic.oup.com/evolut/article/47/1/264/6870150

82. Paradis E. pegas: an R package for population genetics with an integrated–modular approach. Bioinformatics [Internet]. 2010 Feb 1 [cited 2024 Dec 4];26(3):419–20. Available from: 10.1093/bioinformatics/btp696

83. Nei M, Roychoudhury AK. SAMPLING VARIANCES OF HETEROZYGOSITY AND GENETIC DISTANCE. Genetics [Internet]. 1974 Feb 1 [cited 2024 Aug 15];76(2):379–90. Available from: https://academic.oup.com/genetics/article/76/2/379/5990983

84. Akaike H. A new look at the statistical model identification. IEEE Trans Autom Control [Internet]. 1974 Dec [cited 2024 Aug 15];19(6):716–23. Available from: https://ieeexplore.ieee.org/document/1100705

85. Saha A, Kurland S, Kutschera VE, Díez-del-Molino D, Ekman D, Ryman N, et al. Monitoring genome-wide diversity over contemporary time with new indicators applied to Arctic charr populations. Conserv Genet [Internet]. 2024 Jan 20 [cited 2024 Jan 24]; Available from: 10.1007/s10592-023-01586-3

86. Pickrell JK, Pritchard JK. Inference of Population Splits and Mixtures from Genome-Wide Allele Frequency Data. PLOS Genet [Internet]. 2012 Nov 15 [cited 2024 Dec 4];8(11):e1002967. Available from: https://journals.plos.org/plosgenetics/article?id=10.1371/journal.pgen.1002967

87. Auton A, McVean G. Recombination rate estimation in the presence of hotspots. Genome Res [Internet]. 2007 Jan 8 [cited 2024 Oct 28];17(8):1219–27. Available from: https://genome.cshlp.org/content/17/8/1219

88. Kamm JA, Spence JP, Chan J, Song YS. Two-Locus Likelihoods Under Variable Population Size and Fine-Scale Recombination Rate Estimation. Genetics [Internet]. 2016 Jul 1 [cited 2024 Dec 4];203(3):1381–99. Available from: https://academic.oup.com/genetics/article/203/3/1381/6065801

89. Brazier T, Glémin S. Diversity in Recombination Hotspot Characteristics and Gene Structure Shape Fine-Scale Recombination Patterns in Plant Genomes. Mol Biol Evol. 2024 Sep 20;41(9).

90. Gustafsson F, Gunnarsson F. Positioning using time-difference of arrival measurements. In: 2003 IEEE International Conference on Acoustics, Speech, and Signal Processing, 2003 Proceedings (ICASSP ’03). 2003. p. VI–553.

