## Supplementary material for "Leveraging whole genomes, mitochondrial DNA, and haploblocks to decipher complex demographic histories: an example from a broadly admixed arctic fish": Fig. S

Table S1: Sampling sites for Arctic Char in Canada and Greenland

| ID | POP | NAME | REGION | LAT | LON | SAMPLING YEAR | N (>1X) |
| --- | --- | --- | --- | --- | --- | --- | --- |
| 1 | HRN | Hornaday | Inuvialuit Settlement Region, NWT | 69.292090 | -123.748832 | 2011 | 32 |
| 2 | KJA | Kujjuua | Inuvialuit Settlement Region, NWT | 71.203048 | -116.619984 | 2010 | 28 |
| 3 | QNQ | Quunnuq | Inuvialuit Settlement Region, NWT | 69.940459 | -112.621100 | 2010 | 40 |
| 4 | COP | Coppermine | Kitikmeot, Nunavut | 67.810440 | -115.080560 | 2019 | 38 |
| 5 | HAL | Halokvik River | Kitikmeot, Nunavut | 69.162878 | -107.089183 | 2013, 2017 | 36 |
| 6 | EKA | Ekalluk River | Kitikmeot, Nunavut | 69.406836 | -106.316685 | 2014, 2017 | 26 |
| 7 | JAY | Jayko | Kitikmeot, Nunavut | 69.771768 | -103.295213 | 2015, 2017 | 32 |
| 8 | IQG | Iqalungmiut River (Gjoa Haven Fishing Weir) | Kitikmeot, Nunavut | 68.932410 | -96.221350 | 2022 | 40 |
| 9 | NTS | Netsilik Lake | Kitikmeot, Nunavut | 69.432165 | -93.265401 | 2022 | 40 |
| 10 | ABL | Abernethy Lake | Kitikmeot, Nunavut | 71.031961 | -93.393307 | 2022 | 40 |
| 11 | PAM | Pamiurluk Lake | Qikiqtaaluk, Nunavut | 67.083330 | -87.064440 | 2021 | 34 |
| 12 | SAT | Satuut | Qikiqtaaluk, Nunavut | 72.730400 | -80.218900 | 2019 | 39 |
| 13 | KUL | Kuluktoo Bay | Qikiqtaaluk, Nunavut | 72.091400 | -80.934200 | 2019 | 40 |
| 14 | SAM | Sam Ford Fjord | Qikiqtaaluk, Nunavut | 70.791843 | -70.555568 | 2018 | 21 |
| 15 | CFD | Confederation Fiord | Qikiqtaaluk, Nunavut | 68.166667 | -67.316667 | 2017 | 38 |
| 16 | PAD | Paddle Lake | Qikiqtaaluk, Nunavut | 66.816000 | -63.896200 | 2019 | 38 |
| 17 | ISU | Isuituq (PG080) | Qikiqtaaluk, Nunavut | 66.616667 | -67.866667 | 2011 | 40 |
| 18 | QSG | Qasigiat (PG015) | Qikiqtaaluk, Nunavut | 64.616667 | -66.316667 | 2010 | 37 |
| 19 | SVG | Sylvia Grinnell | Qikiqtaaluk, Nunavut | 63.736330 | -68.564941 | 2018 | 28 |
| 20 | KND | Kendall Strait | Qikiqtaaluk, Nunavut | 62.094800 | -65.940359 | 2019 | 38 |
| 21 | QNB | Qinngu, Blandford Bay (LH001) | Qikiqtaaluk, Nunavut | 63.596803 | -71.239397 | 2007 | 36 |
| 22 | KGJ | Kuugarjuk | Kivalliq, Nunavut | 66.471452 | -85.301267 | 2019 | 31 |
| 23 | AKL | Akaalik | Kivalliq, Nunavut | 62.837205 | -91.307115 | 2020 | 30 |
| 24 | CRB | Corbett Inlet | Kivalliq, Nunavut | 62.466667 | -92.333333 | 2020 | 21 |
| 25 | TJL | Ipikituk (IPI) and Saputaliuk (SPI) Rivers | Nunavik, Québec | 58.731000 | -78.375000 | 2018 | 39 |
| 26 | KOA | Korak River | Nunavik, Québec | 60.752623 | -69.792604 | 2018 | 24 |
| 27 | FRM | Francois-Malherbe Lake | Nunavik, Québec | 62.036923 | -74.247532 | 2017 | 35 |
| 28 | PAY | Payne River | Nunavik, Québec | 60.014336 | -70.704358 | 2017 | 27 |
| 29 | CHR | Red Dog River | Nunavik, Québec | 59.297767 | -69.792604 | 2016 | 38 |
| 30 | GEO | George River | Nunavik, Québec | 58.691383 | -65.952957 | 2018 | 34 |
| 31 | UMM | Sermeerlat Kangerluat Lake & Eqaluit Lake | Greenland | 70.540830 | -50.767830 | 2014 | 27 |
| 32 | NUK | Kobbefjord River | Greenland | 64.135180 | -51.383130 | 2013 | 36 |
| 33 | QAQ | Eqaluit Lake | Greenland | 60.761430 | -45.542270 | 2014 | 37 |

Figure S1: Average depth of coverage in populations of the 2X (pop. 1-33) and 8X (three-letter coded), as well as the outgroup population from Sweden. Individuals under 1X (red points) were removed from analyses.

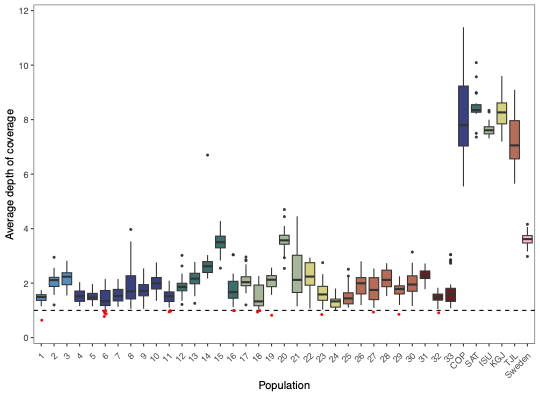

Figure S2: Genome-wide ancestry coefficient for individuals (as vertical bars) in population 1 to 33, as estimated with NGSadmix with number of clusters K = 2, 3, 4, 5, 10, 15, 20, 25, 30, and 35.

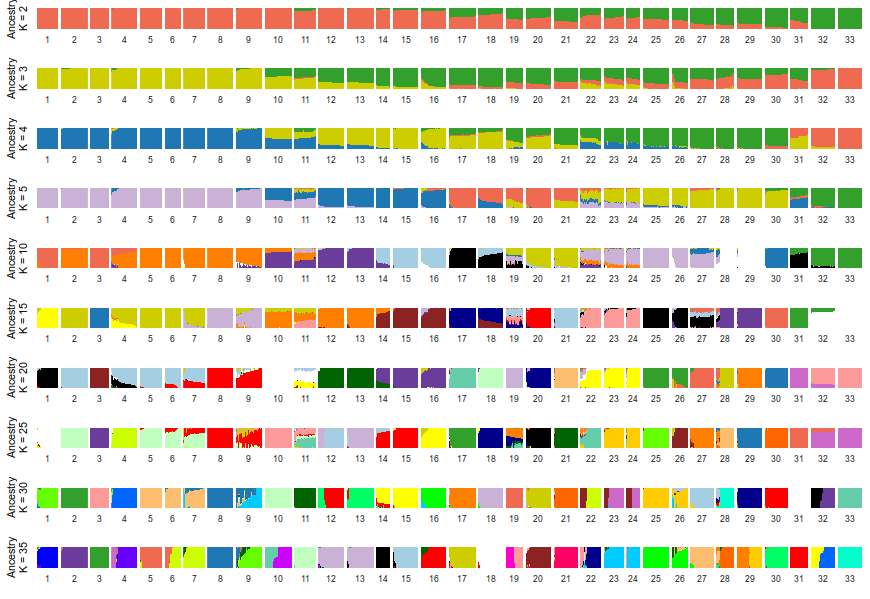
Figure S3: Individual autosomal (genome-wide) heterozygosity per kilobase estimated in 2X data (5X for Sweden, pop. 34).

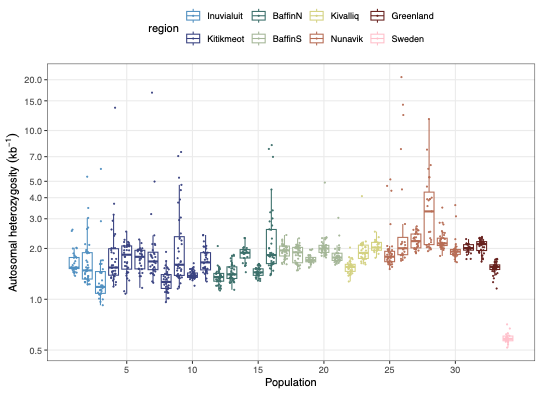

Figure S4: Parameter estimation a) effective size, b) migration and divergence time, and c) effective migration rate in 50 runs of the “ancient gene flow” demographic model in fastsimcoal. See Fig. 5 for the model structure and parameter definition. All scales are logarithmic.

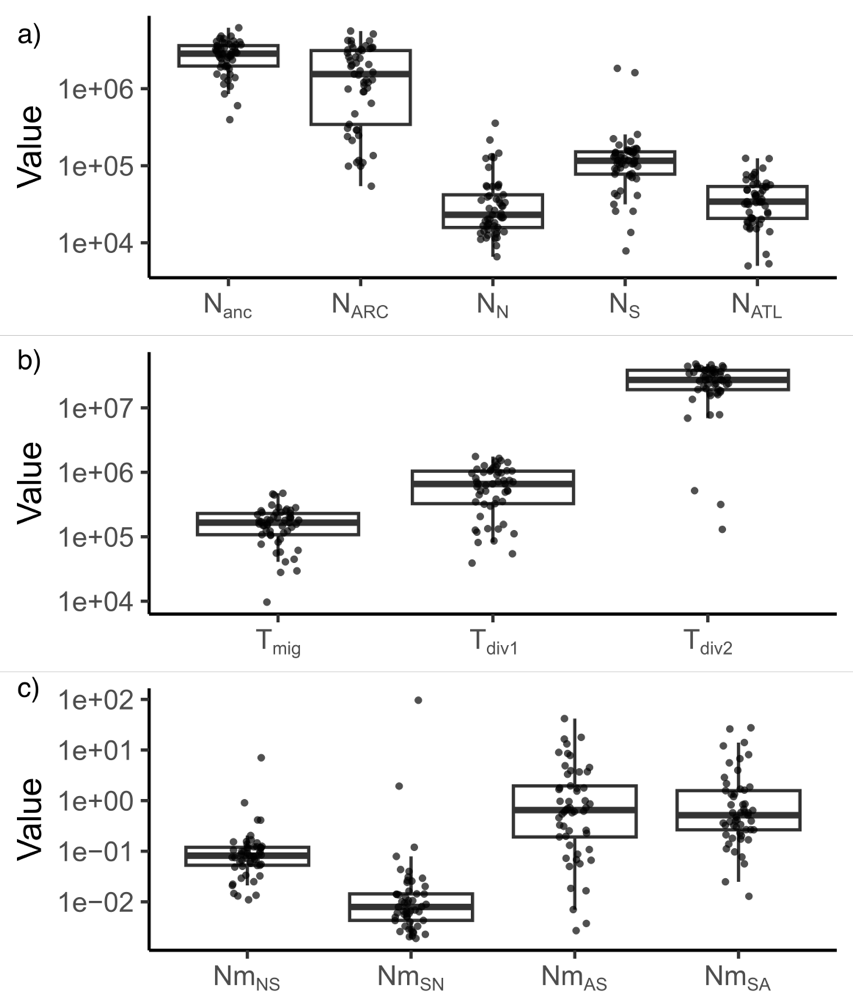

Figure S5: Distribution of genetic diversity by population for the mitochondrial haplotypes of the Arctic lineage along a west-east gradient, approximated by the marine distance from the westernmost population (pop.1). a) Watterson’s estimator and b) nucleotide diversity, standardized by sequence length. c) Tajima’s D. d) Haplotype diversity, i.e. the probability for the randomly selected haplotype in a population to be the same. e) Frequency of the haplotype most observed in the dataset. In every panel, a simple or polynomial (2^nd^) regression curve was added with its 95% confidence of interval, based on the model best supported by AIC.

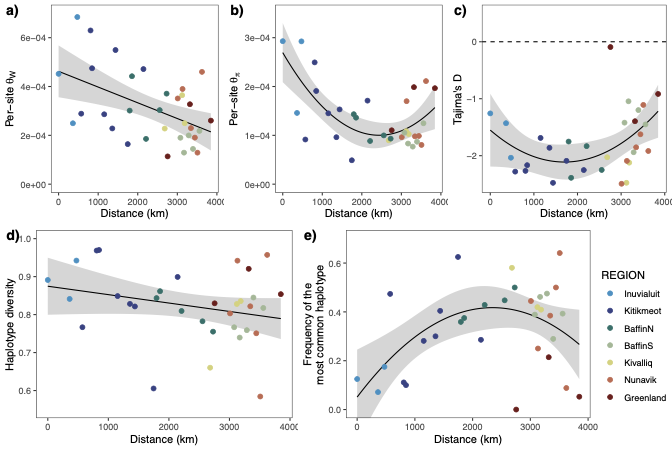

Figure S6: First six multidimensional scaling (MDS) axes summarizing the variation in local PCAs for 100-SNP windows. Every 41 chromosomes (in alternating shades of grey) were analyzed separately. Note that wider gaps are sex-linked chromosomal regions that were excluded from all analyses. Clusters of outlier windows (at least 10 windows over 3 standard deviations away from the chromosomal mean) on the first five axes are noted in red.

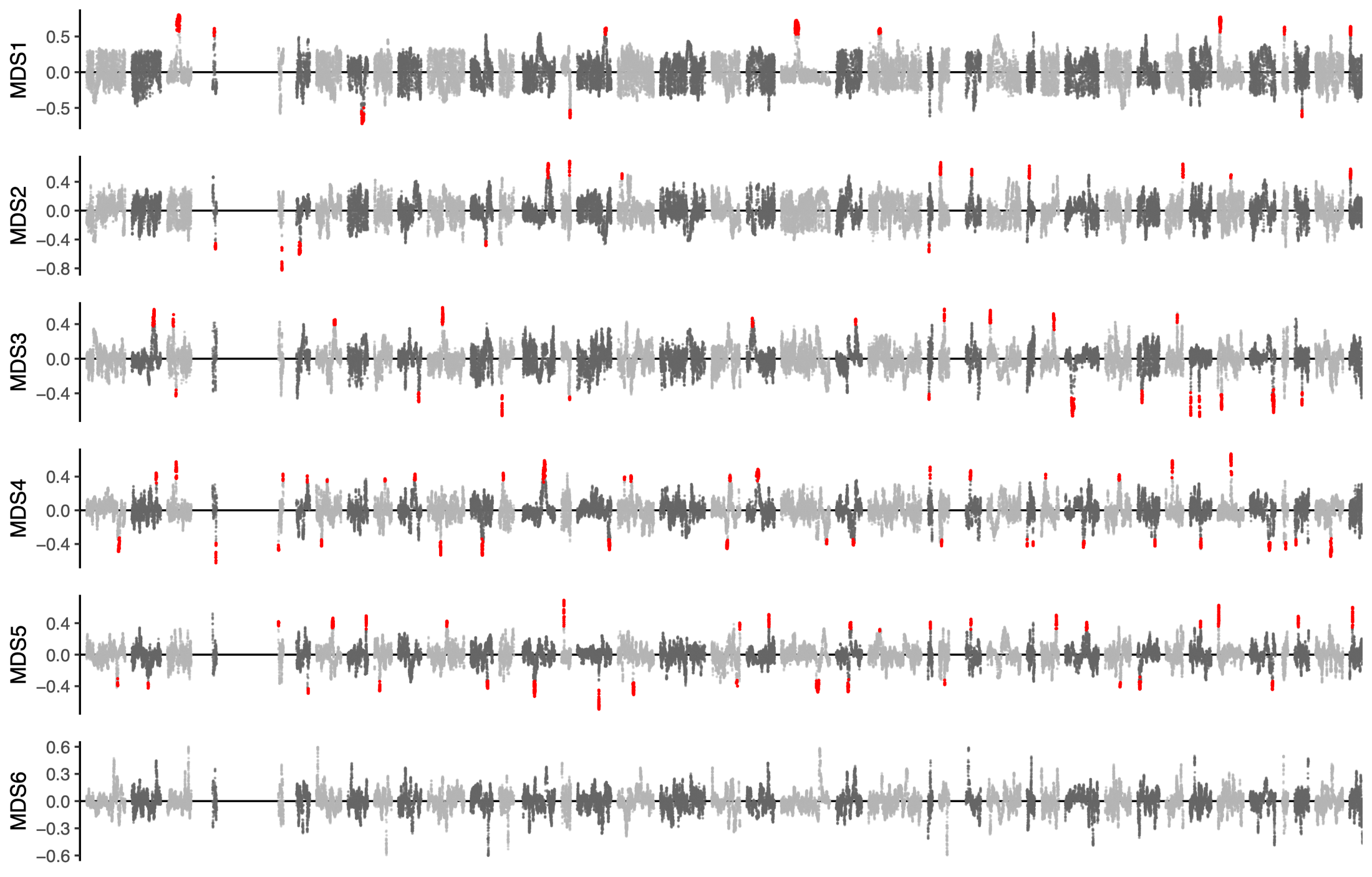

Figure S7: See file **FigureS7.pdf**. All 117 outlier regions identified on the first five MDS axis in a local PCA analysis, as in Fig. 5. a) Position of the MDS outlier region (red) on the linkage group (above) and standardized PC1 coordinates in PCAs for successive non-overlapping 100-SNP windows inside and around the outlier region (below). If three clear clusters were found on PC1, each sample is represented by a line colored according to their haplogroup inferred from panel b). b). PCA for SNPs in the outlier region where the three major clusters found on PC1 are attributed to the 2 homozygotes (AA, blue, left; BB, yellow, right) and their heterozygote (AB, green, middle). Intermediate genotypes (A0, light blue; 0B, light green) were assigned to individuals outside of the cluster distributions. The histogram (above) represents the density of individuals along the first axis of the PCA. c) Proportion of inferred haplogroups for the haploblock across sampling sites.

Figure S8: See file **FigureS8.pdf**. Populational recombination map for each chromosome, estimating the effective recombination rate from 8X data in LDhat, with 95% confidence interval. MDS outlier regions from the local PCA are highlighted in light grey or in light red for the 23 putative haploblocks fitting the Arctic-Atlantic admixture cline described in the main text.

Figure S9: a) Average recombination rate (Rho/kb) in putative local ancestry tracts (haploblock) compared to their chromosome-wide average. Recombination maps were produced for each population in the 8X dataset and for each chromosome separately (see. Fig S9). b) Density plot of the difference in average recombination rate between local ancestry tracts and their chromosome.

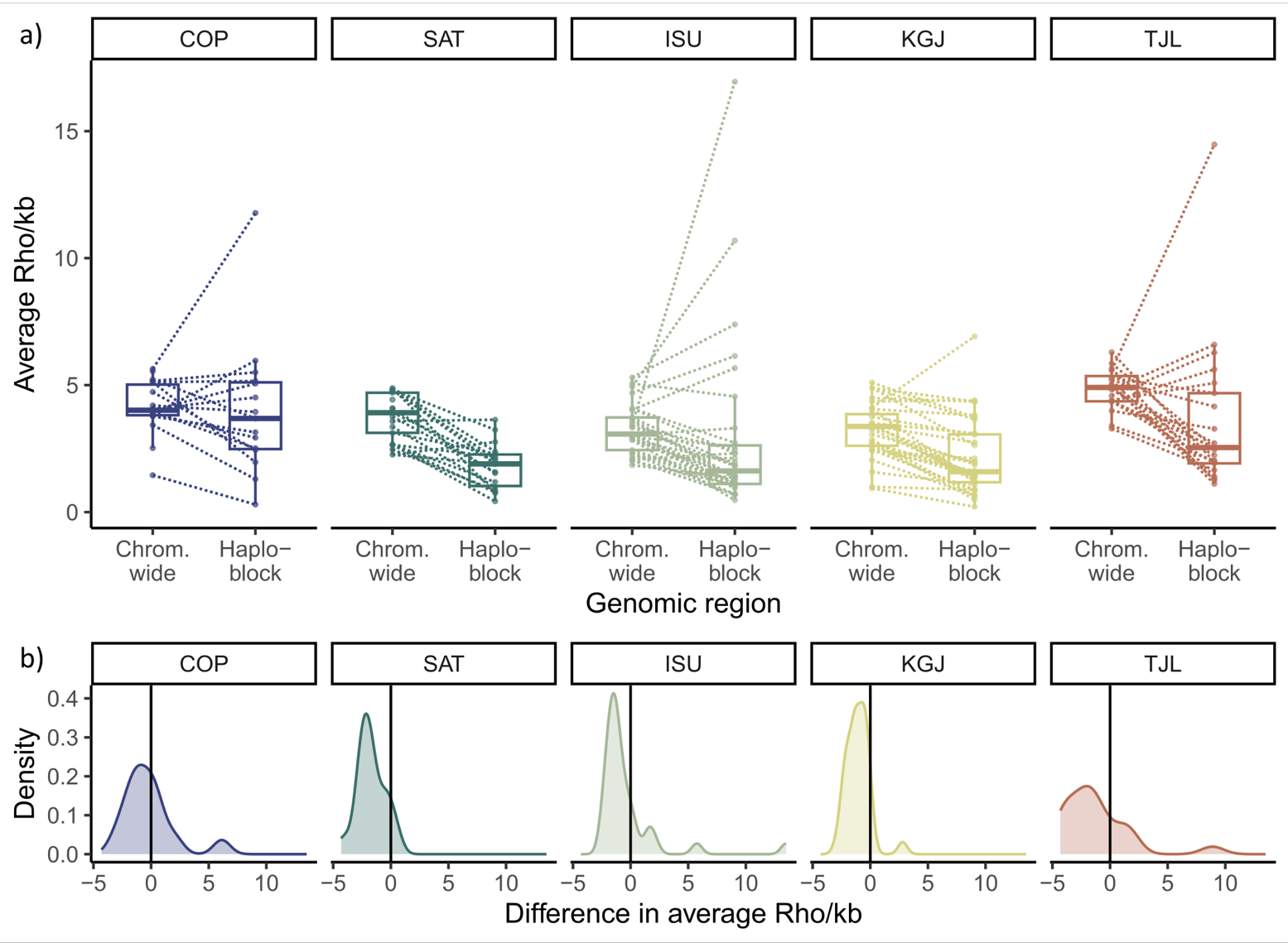

Figure S10: Time Difference of Arrival (TDoA) analysis for inference of the origin of recolonization using directionality indices (ψ) calculated in Arctic haplogroup homozygotes (10 individuals per population) in a) 23 putative local ancestry tracks and b) 23 random genomic regions. Lines indicate rescaled R^2^ = 0.2 isoline for 25 independent runs on different subsets of homozygotes (n = 10) and local ancestry tracts (n = 21). Tile transparency represents the mean rescaled R^2^ over 100 replicate runs as in Fig. 6.

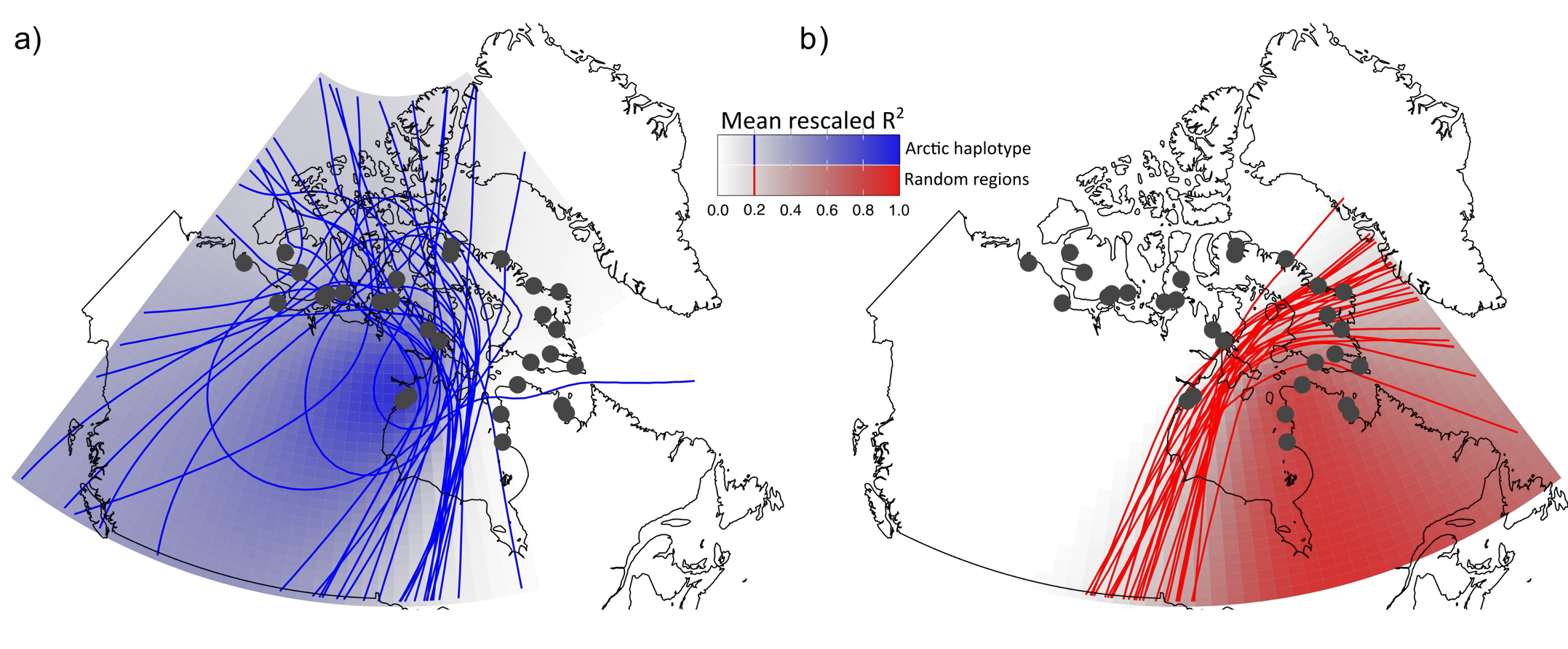
